# Global carbon fixation in Earth’s subsurface biosphere

**DOI:** 10.64898/2026.05.06.723307

**Authors:** Benoit de Pins, Guillermo Climent Gargallo, Martina Cascone, Matteo Selci, Flavia Migliaccio, Deborah Bastoni, Angelina Cordone, Alberto Vitale Brovarone, Stefano Caliro, Gerdhard L. Jessen, J. Maarten de Moor, Peter H. Barry, Karen G. Lloyd, The CoEvolve Project Consortium, Donato Giovannelli

## Abstract

The subsurface of our planet hosts 15% of Earth’s biomass and plays a key role in mediating the exchange of volatiles and elements between deep, long-residence-time geological reservoirs and rapidly cycling surface environments, influencing planetary climate and habitability. While a significant fraction of subsurface microorganisms rely on surface-derived organic carbon, an unknown portion is sustained through chemolithotrophic carbon fixation. Despite its importance, the global diversity and distribution of microbial carbon fixation pathways in the subsurface, and the environmental drivers shaping them, remain poorly constrained. Here we systematically characterise carbon fixation pathways for 412 subsurface metagenomes, including 242 new metagenomes, and compare them to surface-derived datasets. We find that subsurface environments span a broader physicochemical space than surface systems and support a higher abundance and diversity of carbon fixation strategies. Using colocated geochemical data spanning >50 variables, we show that the reductive tricarboxylic acid cycle and the reductive acetyl-CoA pathway are enriched in reducing, geochemically evolved fluids. We use the metagenomic results together with previously published carbon fixation rates in the subsurface to derive a global continental subsurface carbon fixation rate of ∼2.65 Pg C yr^−1^ (range: 0.31–2.99). This represents ∼2% of terrestrial photosynthetic primary production, and is an order of magnitude higher than geological fluxes between the surface and the subsurface. These results identify the subsurface as a reservoir of autotrophic strategies organized along geochemical gradients, contributing substantially to the global carbon cycle.

**One Sentence Summary:** The subsurface is a widespread, environmentally and functionally diverse reservoir of autotrophic carbon fixation pathways that can contribute substantially to the global carbon cycle, fixing ∼2.65 Pg C yr^−1^ in continental settings alone.

## Main

Carbon fixation, the process by which inorganic carbon enters the biosphere, underpins all food webs on Earth. This process is carried out by autotrophs, organisms found across the three domains of life, which have evolved multiple metabolic strategies to assimilate inorganic carbon under distinct physicochemical constraints. To date, at least seven carbon fixation pathways have been described: the Calvin Benson Bassham (CBB) cycle found in plants, algae, and many aerobic and microaerophilic bacteria; the Wood–Ljungdahl (WL) pathway (also known as the reductive acetyl-CoA pathway) found in acetogenic and sulfate-reducing bacteria, and methanogenic and sulfate-reducing archaea; the reductive glycine pathway found in some anaerobic bacteria and archaea; the reductive tricarboxylic acid (rTCA) cycle (also known as the Arnon-Buchanan cycle) used by anaerobic and facultative microaerophilic bacteria and some thermophilic archaea; and three strongly overlapping cycles (the 3-hydroxypropionate/4-hydroxybutyrate (HPHB) and dicarboxylate/4-hydroxybutyrate (DCHB) cycles, found in archaea; and the 3-hydroxypropionate (3HP) found in some bacteria)^1^. These pathways differ markedly in their energetic efficiency, oxygen sensitivity, and cofactor requirements, suggesting that their distribution is environmentally constrained^2,3^. While carbon fixation at Earth’s surface mostly relies on oxygenic photosynthesis and the CBB cycle, fixation in the deep biosphere relies on alternative pathways adapted to distinct geochemical conditions^4,5^.

Subsurface ecosystems are defined as the portion of lithosphere between the near surface of our planet (one to a few metres depth in the soil or marine sediments) and the depth of the 122-135 °C isotherm representing the current boundary of known life forms^6^. Containing up to ∼15% of Earth biomass, subsurface ecosystems host approximately 7 × 10^29^ microbial cells, constituting around 60% of all bacteria and archaea on Earth^7^. These ecosystems can be largely decoupled from photosynthetic inputs and are instead sustained by geochemical energy sources, supporting widespread chemoautotrophy^8,5^. Subsurface microbial communities have been shown to influence long-term carbon cycling by mediating the removal of large fractions of carbon cycled through geological processes (e.g., subduction and volcanism)^8–11^.

While individual studies have documented the diversity of carbon fixation pathways in specific subsurface settings^4,5,12–14^, a comprehensive, functionally resolved view across global subsurface ecosystems is lacking. In particular, it remains unclear whether carbon fixation strategies are distributed homogeneously across sites, or whether they follow consistent organisational patterns structured by environmental gradients, and how these patterns scale to constrain the global contribution of subsurface chemolithotrophy to the carbon cycle. We analysed more than 400 subsurface metagenomes alongside hundreds of publicly available surface datasets, and integrated these with environmental data to identify the main determinants of *in situ* carbon fixation pathways. We show that subsurface carbon fixation genes are consistently more abundant and diverse than in the surface, and are organized into coherent functional regimes associated with geochemical gradients, particularly temperature and redox conditions. By linking genomic potential to fluid-derived volumetric rates, we further constrain the contribution of continental subsurface chemolithotrophic carbon fixation to the global carbon cycle.

### Subsurface environments span a wide physicochemical space

The deep subsurface is the most genetically diverse ecosystem on Earth, encompassing the majority of uncultured bacteria and archaea^15^. However, access to these environments has traditionally relied on deep drilling projects, which are expensive and infrequent^16^. Natural seeping fluids represent an alternative window to subsurface ecosystems^17,18^. We collected sediments, fluids, and occasionally biofilms from 176 deeply-sourced seeps worldwide, combining *in situ* physicochemical measurements with their metagenomic sequencing and a characterization of their geochemistry (see Materials and Methods), resulting in 373 metagenomes, of which 330 passed quality filtering. These data are complemented by 82 publicly available subsurface metagenomes from the literature. We compared these to 164 oceanic metagenomes from the Tara Oceans and Malaspina expeditions^19,20^, more than 500 freshwater metagenomes, and >350 soil metagenomes from the National Ecological Observatory Network (NEON; see Materials and Methods), supplemented with an additional 41 publicly available soil and freshwater metagenomes from the literature. Together, these datasets of 1,513 samples span a broad geographic, tectonic, geochemical and environmental space (Fig. 1a and Supplementary Fig. 1). Analysis of pH and temperature distributions (Fig. 1b), water chemistry (Fig. 1c), and additional physicochemical parameters (Supplementary Fig. 2), shows that subsurface environments occupy a broader and more heterogeneous physicochemical space than surface ecosystems. They span a larger fraction of the global pH–temperature space and exhibit greater hydrochemical diversity (pH–temperature convex hull: 86% of normalised space vs. ≤43% for all other groups; Piper diamond coverage: 82% vs. 45%; hydrochemical facies diversity: Shannon H = 1.90 vs. 0.93 bits). This broader physicochemical diversity likely reflects the combined effects of lithological heterogeneity and prolonged water-rock interaction, which generate a wide range of redox states and fluid chemistries in the subsurface. Unlike surface environments, which are rapidly homogenized by mixing and atmospheric exchange, subsurface systems evolve over long residence times and maintain geochemically isolated regimes, allowing metabolic niches to diverge and persist.

**Figure 1.**
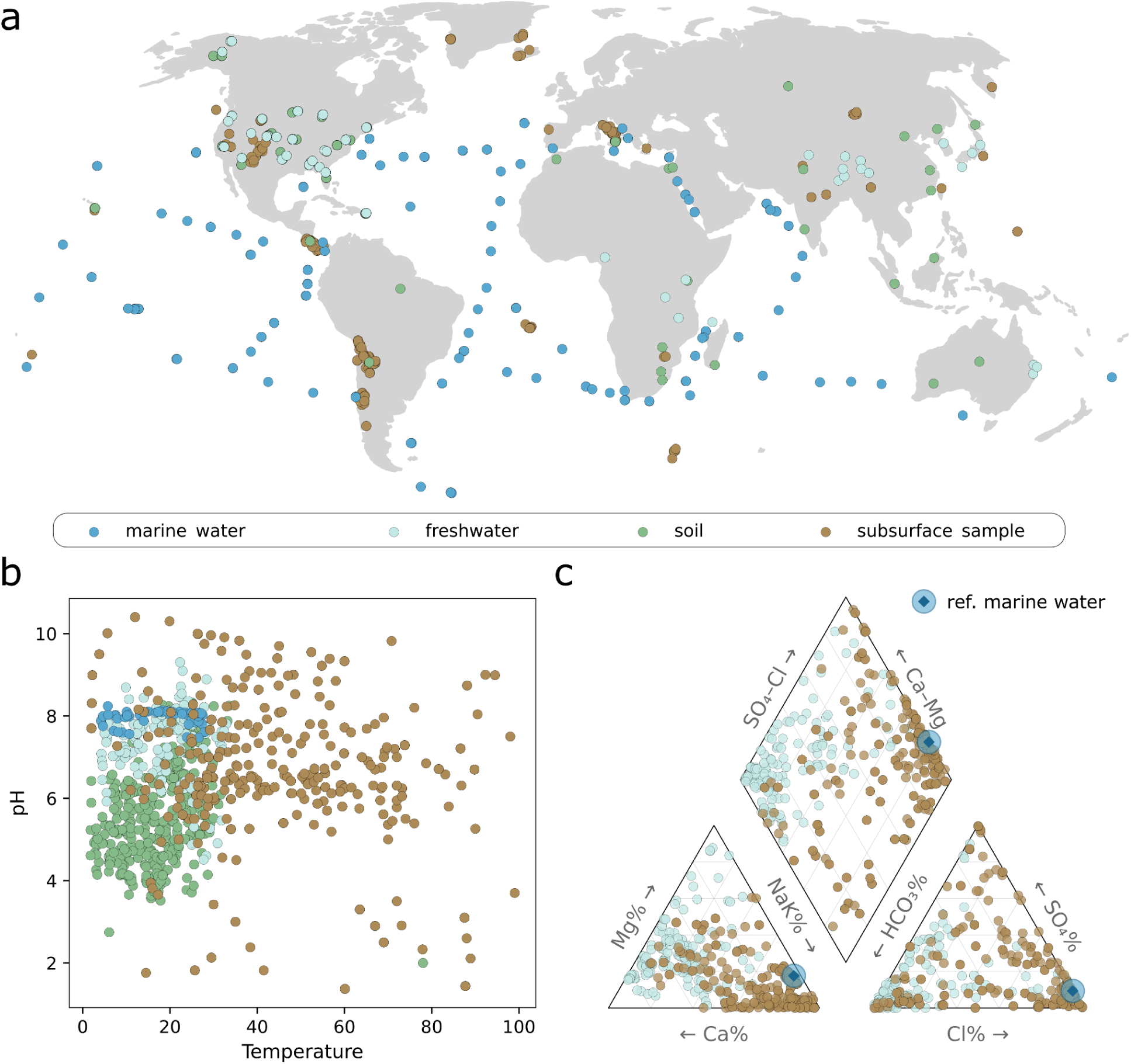
Global distribution and physicochemical space of subsurface and surface metagenomes. **a**, Global map of all metagenomic sampling sites included in this study, comprising geothermal and subsurface fluids, sediments, and biofilms from the CoEvolve dataset, as well as publicly available surface (soil, marine, freshwater) and subsurface samples from public repositories. **b,** Distribution of samples in pH–temperature space, highlighting the substantially broader physicochemical range covered by subsurface environments compared to surface ecosystems. **c,** Piper diagram summarising aqueous geochemistry for samples with available data. The lower triangles represent the relative proportions of major cations (Ca^2+^, Mg^2+^, Na^+^+K^+^) and anions (Cl^−^, SO_4_^2-^, HCO_3_^−^), while the central diamond integrates these to define hydrochemical facies. Seawater samples are represented by a narrow area (blue diamond circled). Soil and sediment samples are absent from this plot.

### Subsurface microbial communities occupy a distinct global functional space

We applied a single, unified pipeline to functionally annotate all 1,513 metagenomes (see Materials and Methods). Briefly, for each metagenome, we quantified the abundance of each KEGG Ortholog (KO) and normalised it to the β subunit of RNA polymerase, accounting for both bacterial and archaeal homologs. Our analysis, performed at the read level, enables the detection of low-abundance functions that may be lost in assembly-based methods, and therefore permits consistent comparisons across large datasets^21^. This allowed for the identification and comparison of carbon fixation pathways across hundreds of metagenomes, revealing global trends. Multivariate analysis of global KO composition reveals clear clustering by environment type (Fig. 2a). Metagenomes originating from similar environments mostly clustered together regardless of their source (*e.g.*, subsurface metagenomes from this study and those retrieved from the literature, see Supplementary Fig. 3), indicating no dataset driven batch effects. Functional composition differed strongly among environments (PERMANOVA, F = 178.1, R^2^ = 0.26, p ≤ 0.001), and multivariate dispersion varies significantly between groups (PERMDISP, F = 76.6, p ≤ 0.001). Subsurface communities showed the highest distances to their group centroid (Fig. 2b), indicating a greater functional heterogeneity compared to marine, soil, and freshwater ecosystems.

**Figure 2.**
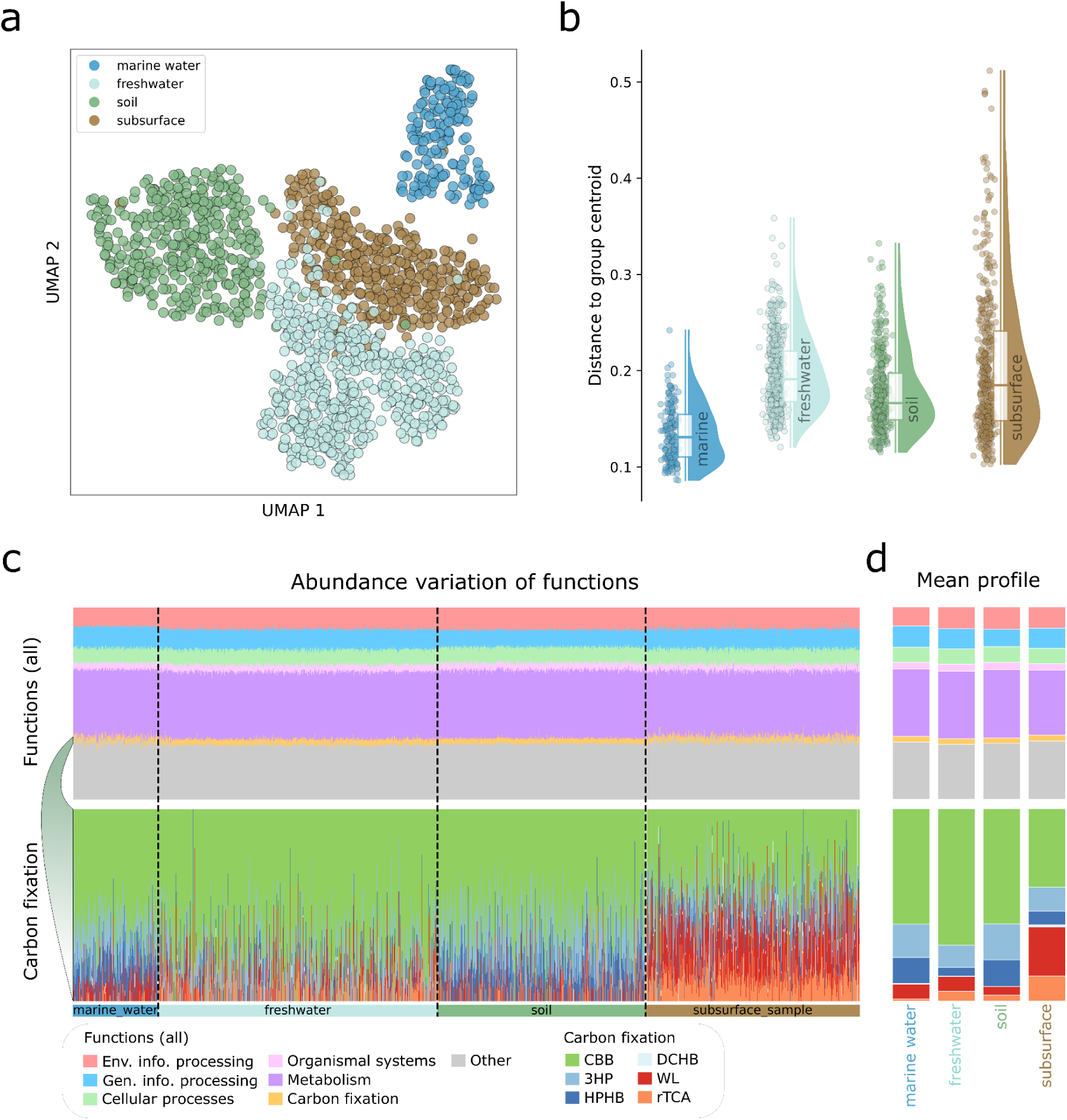
Functional differentiation of subsurface and surface microbial communities. **a**, Uniform Manifold Approximation and Projection (UMAP) of all samples based on metagenome-derived functional profiles (KEGG Ortholog abundances), showing a clear separation of subsurface communities from surface marine, soil, and freshwater ecosystems. Points represent individual samples and are coloured by environment. **b,** Quantification of functional separation among environments: distances of individual samples to their environment-specific centroid in the original functional distance space, highlighting both the distinctness and internal heterogeneity of subsurface communities. **c,** Relative distribution of broad functional categories across environments, showing substantial redundancy at coarse metabolic levels (e.g., metabolism, information processing), contrasted with pronounced differences in carbon fixation functions. KOs annotated to multiple functional categories were counted in each respective category; relative abundances represent the proportional functional signal per category and do not sum to 1. **d,** Mean functional profiles across environments.

### Functional separation persists despite broad metabolic redundancy

Despite clear separation in multivariate space (Fig. 2a), global functional profiles remained broadly similar across environments (Fig. 2c), consistent with previous observations of metabolic redundancy at the ecosystem scale^22^. This indicates that biome differentiation is not driven by global metabolic capacity, but by the differential organization of specific functional modules in diverse environments. Carbon fixation represents one such module, showing strong and systematic divergence between surface and subsurface environments (Fig. 2c, bottom panel). Indeed, carbon fixation KO abundances show marked differences across all environments, with subsurface ecosystems displaying the most distinctive profiles among all environments analysed. Subsurface environments represent oligotrophic niches, poor in organic matter relative to surface ecosystems, probably selecting for a diversity of autotrophic lifestyles^23^.

### Carbon fixation potential is enriched and diversified in the subsurface

To quantify the abundance of each carbon fixation pathway without conflating it with genes shared with other central metabolic functions, we compiled a list of pathway-specific marker genes (see Supplementary Note 1, Supplementary Table 1, and Supplementary Fig. 4 for an analysis showing why global KO-level comparisons fail to capture this signal). Since the reductive glycine pathway overlaps with the WL pathway in several key enzymes, we combine them as a single WL-type module. Consistent with the pattern observed in Fig. 2c and d, subsurface communities exhibit both higher abundance and greater diversity of carbon fixation pathways (Fig. 3a; Mann–Whitney U test, p < 0.001, Supplementary Fig. 5; Mann–Whitney U test, p < 0.001). These effects are independent of any batch-specific sequencing depth (Supplementary Note 2 and Supplementary Fig. 6). The observed increased abundance and diversity of autotrophic pathways in the subsurface is consistent with adaptation to the nutrient-limited conditions typical of subsurface environments, which likely select for metabolically self-sufficient lifestyles.

**Figure 3.**
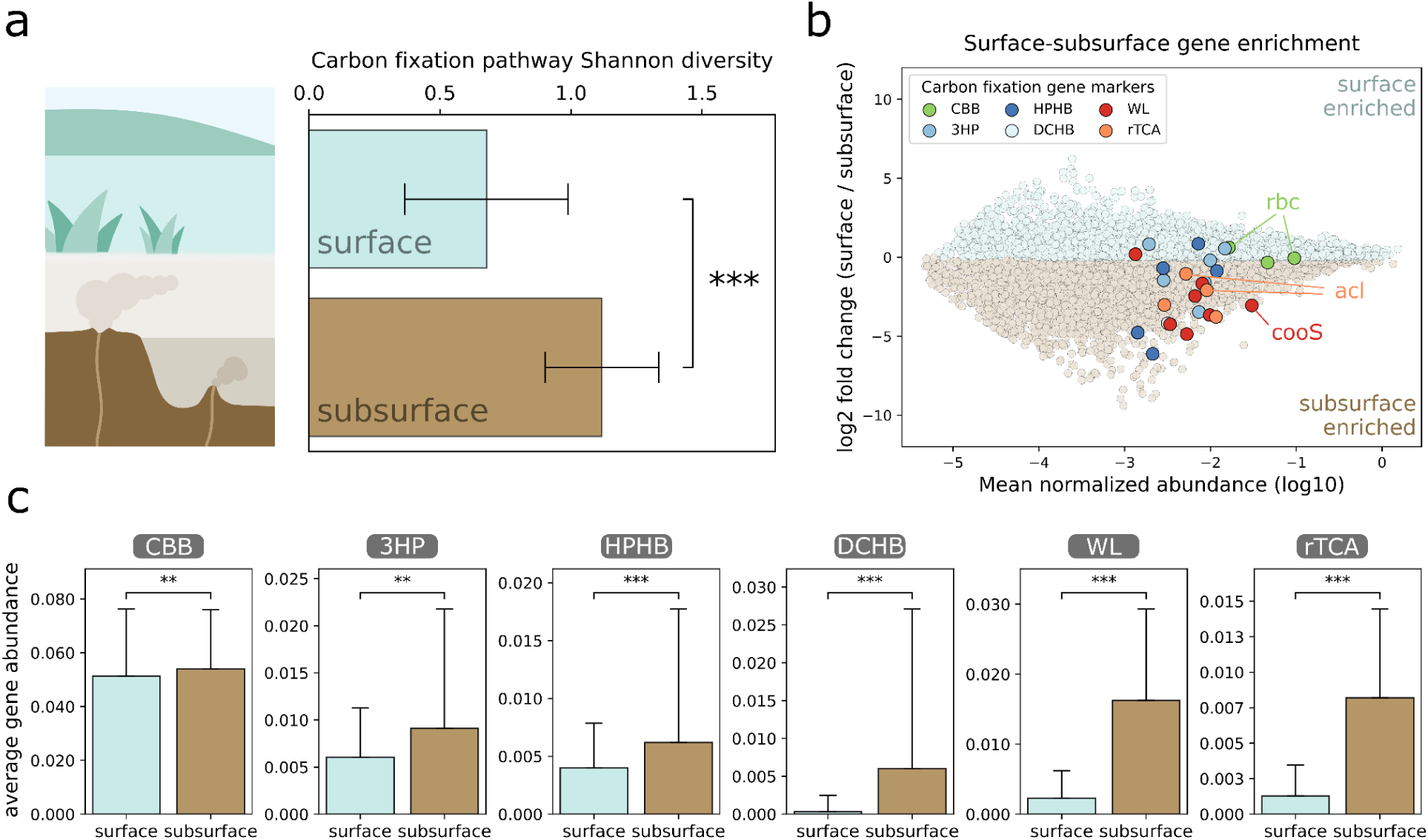
Carbon fixation potential is enriched and diversified in subsurface environments. **a**, Comparison of total carbon fixation gene diversity between surface and subsurface environments, showing a significant enrichment of carbon fixation potential in subsurface samples (Mann-Whitney U test, p < 0.001). **b,** Differential enrichment analysis of individual genes between subsurface and surface samples, plotted as log fold-change versus mean abundance (normalised by rpoB). Carbon fixation gene markers are highlighted. CBB cycle marker genes show limited enrichment, whereas genes associated with alternative carbon fixation pathways, including the rTCA cycle and the WL pathway, are strongly enriched in subsurface environments. **c,** Pathway-level comparison of carbon fixation strategies, showing marked enrichment of pathway genes in the subsurface. Mann-Whitney U test, ** p < 0.01, *** p < 0.001.

When comparing gene abundances between surface and subsurface metagenomes at the global level, most marker genes associated with carbon fixation pathways were among those enriched in subsurface communities (Fig. 3b). A pathway-level comparison confirmed this pattern: all carbon fixation pathways showed higher relative abundance in subsurface environments (Fig. 3c and Supplementary Fig. 7 for a detail of each sample type). The strongest enrichment was observed for the WL pathway and the rTCA cycle, two pathways adapted to low-oxygen conditions and frequently reported in subsurface settings including hydrothermal vents^24^, boreholes^25^, groundwater^26^, hot springs^5^, and cold seeps^27^. On the other hand, the weakest enrichment was observed for the 3HP, the HPHB, and the CBB cycles, which are the three oxygen-tolerant carbon fixation pathways. Together, these results indicate that subsurface communities show a higher abundance and a greater diversity of carbon fixation strategies. This enrichment is mostly due to pathways other than the CBB cycle, suggesting that subsurface ecosystems favour non-canonical pathways adapted to reducing conditions.

### Environmental gradients structure subsurface carbon fixation modules

Network analysis of carbon fixation gene co-occurrence patterns shows that carbon fixation genes in the subsurface are more modular than in surface systems, forming tightly connected functional assemblages (Fig. 4a,b and Supplementary Fig. 8 for a comparison of the network topology for the surface and the subsurface). This increased modularity, as supported by Fig. 3, reflects stronger co-variation of gene abundances across samples, suggesting a constrained environmental control along geochemical gradients in the subsurface. In contrast, surface datasets display more diffuse and locally structured networks, likely reflecting a control by a higher number of controlling environmental parameters (e.g., dispersion, mixing, diel cycles, variable organic carbon inputs, etc.).

**Figure 4.**
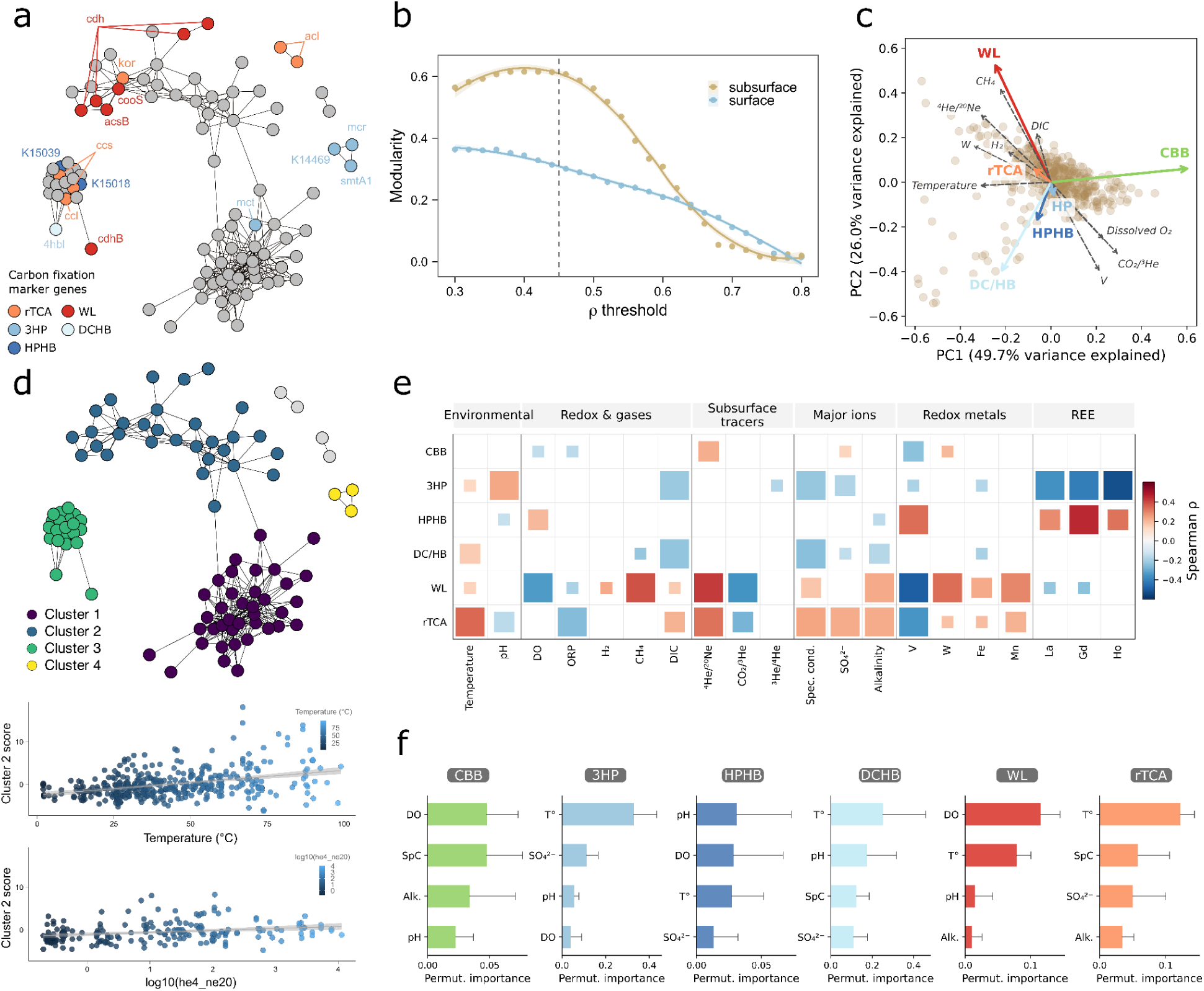
Environmental structure and drivers of carbon fixation pathways in the subsurface. **a**, Subsurface carbon fixation gene co-occurrence network at ρ=0.45. **b,** Network modularity across correlation thresholds for surface and subsurface metagenomes. **c,** Principal component analysis (PCA) of the abundances of the six carbon fixation pathways across subsurface samples (n = 410). Each point represents a sample. Coloured arrows indicate pathway loadings in ordination space. Selected environmental variables significantly associated with pathway composition are shown as dashed vectors. **d,** Louvain clusters derived from the subsurface carbon fixation gene co-occurrence network at ρ = 0.45 (top), and correlations of cluster 2 with temperature and log_10_(^4^He/^20^Ne) (bottom). **e,** Spearman correlations between carbon fixation pathways and environmental variables measured in collected samples. Square size represents statistical significance (FDR–BH q-value) and colour indicates the correlation coefficient (ρ). Variables are grouped into general environmental parameters, redox and gas indicators, subsurface tracers, major ions, redox-sensitive metals, and rare earth elements (REE). **f,** Random Forest permutation importance for the prediction of carbon fixation pathways abundances from 6 high-coverage physicochemical variables. Importance scores are the mean decrease in R^2^ upon variable permutation (5-fold CV); error bars show the standard deviation across folds.

To identify the possible drivers of this organisation in the subsurface, we integrated metagenomic data with >50 environmental variables measured in the field or on co-located, synoptically collected samples for the subsurface dataset (or collected from published data when available for the surface samples). These variables encompass physico- and geochemical parameters, major ions, trace metals, reactive volatiles and noble gases (see Materials and Methods). The ordination of the pathway abundance profiles revealed a separation of the different carbon fixation pathways along dominant environmental axes (CBB in one; WL and rTCA in another; 3HP, DCHB, and HPHB in another). It also revealed the principal parameters associated with these profiles, including temperature, redox-sensitive metals (vanadium and tungsten shown here), crustal tracers, and several gases (Fig. 4c and Supplementary Fig. 9). These different axes capture coupled gradients in temperature, redox state, and fluid evolution, and therefore indicate that the distribution of the different carbon fixation pathways reflects underlying geochemical regimes rather than contingencies specific to single sites.

Among the different pathways, the rTCA cycle and the WL pathway show consistent environmental associations, being systematically enriched in geochemically evolved, reducing fluids. These fluids are characterised by elevated ^4^He/^20^Ne and low CO_2_/^3^He, indicating deeply sourced fluids (environmental vector fitting in Fig. 4c, confirmed by pairwise Spearman correlations in Fig. 4e). Moreover, the rTCA cycle is strongly associated with high temperatures, in accordance with its observed high prevalence in thermophilic and hydrothermal environments^28,29^ while the WL pathway shows the highest anti-correlation with dissolved oxygen among all pathways, consistent with the strict anaerobic requirement of its key enzymes^30,31^. The WL pathway is also correlated with elevated H_2_ and CH_4_, reflecting its coupling to hydrogenotrophy and methanogenesis^31^. Trace metals provide additional signatures of these environments^32^: while key enzymes of the WL pathway, like the formate dehydrogenase, can incorporate either molybdenum or tungsten as cofactors depending on the organism^33^, the abundance of the WL pathway is correlated to tungsten levels but not to molybdenum in the subsurface (Supplementary Fig. 10 for a complete picture of correlation with all environmental factors), suggesting that the highly reducing conditions of these environments promote the selection of W-dependent variants. Supporting this, while in reducing, sulfide-rich and hydrothermal fluids, Mo can be depleted by precipitation as molybdenite, or by conversion into thiomolybdates (which are subsequently incorporated into sulfide-associated phases), W remains comparatively more soluble and enriched in evolved geothermal fluids^34^. Accordingly, W-dependent formate dehydrogenases and other tungstoenzymes are widespread in anaerobic and thermophilic microorganisms, where they catalyse oxygen-sensitive, low-potential redox reactions^35,36^. The preferential association of WL marker genes with W therefore suggests that not only subsurface environments select for this carbon fixation pathway, but also for the metal-cofactor variants that are best adapted to trace metals availability in these ecosystems. Convergent evidence from the co-occurrence network, where several marker genes of the WL pathway together with rTCA are gathered in a single cluster whose abundance correlates with temperature and ^4^He/^20^Ne across samples (Fig. 4d), and from random forest regression, predicting pathway abundance from environmental variables, which identifies temperature and dissolved oxygen as the main predictors of the rTCA cycle and the WL pathway respectively (Fig. 4f), confirm that these two pathways occupy largely overlapping geochemical niches, defined by deep, hot, and reducing conditions, in accordance with their enzymatic constraints and known energetic efficiencies.

The other carbon fixation pathways were associated with distinct environmental factors, revealing the multidimensional nature of the environmental control of subsurface carbon fixation. 3HP, HPHB, and DCHB pathways seem associated with more diluted and surface-influenced fluids, temperature remaining a predictive factor but in a distinct geochemical context (Fig. 4e,f). The CBB cycle showed only few and weak correlations to environmental parameters (Fig. 4e) and a significant but weak predictability by the random forest model (R^2^ = 0.08, p ≤ 0.001, Fig. 4f). This probably reflects its large ecological distribution across various environments, as observed here and reported in previous studies^5,37,38^.

To link the distribution of the different carbon fixation pathways to an ecosystem potential, we defined a Carbon Fixation Potential Index (CFPI), as the sum of rpoB-normalised pathway scores. CFPI reflects the autotrophic capacity of a metagenome (see Materials and Methods). It correlates with temperature (ρ = 0.39), ^4^He/^20^Ne ratios (ρ = 0.37), and negatively correlates with dissolved oxygen (ρ = -0.30; Supplementary Fig. 11a). We also observe opposite trends for tungsten and vanadium. CFPI increases with tungsten (ρ = 0.37) and decreases with vanadium (ρ = -0.37), which fits their contrasting behaviour in reducing environments. Indeed, tungsten shows a higher mobility in sulfidic and hydrothermal fluids due to the solubility of thiotungstate complexes and restricted adsorption on iron oxyhydroxides, while vanadium, usually present as dissolved vanadate in oxygenated surface waters, is reduced and less mobile in reduced environments^39^. The convergence of these physical, isotopic, and geochemical indicators suggest an organisation of carbon fixation across a continuous gradient from oxygenated surface waters to reduced deep geo-fluids.

Principal component analysis of environmental variables also captures this dominant environmental gradient linking the redox state, temperature and geochemical fluid evolution (Supplementary Fig. 11b). CFPI increases along this gradient, confirming that the carbon fixation potential of a subsurface fluid emerges from the combined effects of thermal regime, fluid evolution, and the availability of geochemically derived reducing power.

### Carbon fixation potential of continental subsurface biosphere

To estimate subsurface carbon fixation, we anchored CFPI values to published empirically measured rates in the subsurface and combined them with a geophysically derived estimate of habitable crustal volume. The depth of the 135 °C isotherm was derived from a global heat-flow field^40^, with constraints applied to avoid unrealistic penetration of conductive thermal profiles. The resulting habitable domain reflects the distribution of thermal regimes at the planetary scale, rather than lithological weighting. Integrating across all grid cells, the habitable crust above the 135 °C isotherm spans ∼1.76 × 10^9^ km^3^ of rock and contains ∼6.7 × 10^7^ km^3^ of pore water. This pore water reservoir is partitioned into continental (∼3.6 × 10^7^ km^3^) and oceanic (∼3.1 × 10^7^ km^3^) domains, corresponding to roughly 3–5% of the global ocean volume. These values fall within previously reported constraints on subsurface fluid reservoirs^41–43^. Uncertainty (∼2-3x) is mainly driven by effective porosity and spatial variability in heat flow, particularly in low heat-flow regions where conductive models predict deeper isotherms. By explicitly constraining the depth and porosity, this approach provides a conservative and physically consistent estimate of the volume of subsurface fluids available to support microbial activity.

Since most subsurface samples in this study originate from continental settings, we used the continental pore water volume to derive a metagenomic-informed global estimate of subsurface carbon fixation. CFPI was scaled against volumetric carbon fixation rates constrained by measured subsurface systems^44,4,45,46^, allowing us to obtain a metagenomic weighted distribution of expected carbon fixation rates to extrapolate across heterogeneous environments without assuming uniform activity. This yields a continental subsurface chemolithotrophic carbon fixation rate of 2.65 Pg C yr^−1^ (range: 0.31-2.99 Pg C yr^−1^). This estimate exceeds the conservative 0.11 Pg C yr^−1^ carbonate-groundwater value reported by Overholt et al.^4^, and falls within recent estimates for marine chemosynthesis (1.2-11 Pg C yr^−1^)^47^. While representing ∼2% of terrestrial gross primary production (123 ± 8 Pg C yr^−1^)^48^, this subsurface carbon fixation rate must be contextualized to geological fluxes; it is an order of magnitude higher than the estimated 0.2-0.3 Pg C yr^−1^ volcanic flux to the surface^49,50^. While a substantial fraction of this fixed carbon is likely recycled locally through heterotrophic turnover (for comparison, ∼1.73 Pg C yr^−1^ is released by heterotrophic organic carbon degradation in marine sediments alone^51^), the magnitude of the flux requires a persistent input of inorganic carbon from geological sources, including magmatic degassing and water–rock interactions. In the absence of subsurface chemolithotrophy, a large fraction of the geological carbon would remain mobile and potentially reach the surface, with geological emissions to the atmosphere up to ten times higher. The deep subsurface biosphere therefore operates as a major biological sink on geological carbon fluxes, intercepting and transforming geological carbon before it reaches the atmosphere, mediating the exchange between deep Earth processes and the surface carbon cycle.

## Conclusions

Carbon fixation is the primary gateway through which inorganic carbon enters the biosphere, sustaining all ecosystems. While surface environments are largely dominated by the CBB cycle, the absence of light and diversity of energy sources in the subsurface selects for a broader spectrum of chemolithotrophic pathways. By combining 412 subsurface metagenomes with surface datasets, we show that subsurface carbon fixation is both more abundant and more diverse than at the surface, and is structured along geochemical gradients. In particular, the rTCA cycle and the WL pathway are consistently enriched and associated with reducing, geochemically evolved fluids of deep origin. This indicates that subsurface autotrophy operates as a distinct, geochemically constrained regime adapted to anoxic, energy-limited environments. We note that the boundary between surface and subsurface is not discrete. Low-oxygen soils, sediments, and deep aquifers likely occupy transitional states along the same continuum. Resolving these transitions will be critical to define where and how the organizational logic of subsurface carbon fixation emerges.

When scaled to the continental subsurface, this structure translates into a flux on the order of ∼2.65 Pg C yr^−1^. This magnitude exceeds estimated volcanic carbon inputs to the surface, indicating that a substantial fraction of geologically sourced carbon is intercepted and transformed before it can escape to the atmosphere. Subsurface chemolithotrophy therefore acts as a sink on deep carbon fluxes, coupling geological processes to the surface carbon cycle.

Our estimates remain conservative and first-order. They do not resolve the partitioning between newly fixed carbon and locally recycled carbon, nor capture the full heterogeneity of permeability and porosity at depth. However, they establish a quantitative baseline, revealing that subsurface chemolithotrophy operates at a scale relevant to the global carbon cycle, and the deep biosphere must be considered an active component of planetary carbon budgets.

## Materials and Methods

### Sample collection and *in situ* measurements

Sampling was performed in the frame of the ERC funded project CoEvolve in globally distributed locations representing diverse geological settings (Supplementary Fig. 1, CoEvolve samples), and following a large-scale sampling approach described previously^18^. Where possible, for each location three types of samples were collected: fluids, sediments, and biofilms. The full sampling procedure is described in detail elsewhere^52^.

Fluid samples (500 mL to 3 L, depending on whether the filter clogged from precipitates) were collected from the main seeping point of the sampled feature by inserting a titanium tube connected to a silicone tube into the orifice, when possible. Otherwise, fluids were collected as freshly-expressed as possible. Fluids were collected using a portable peristaltic pump or 60 mL syringes and directly filtered through multiple 0.22 μm polyethersulfone cartridge filters (Sterivex, Millipore Sigma), and stored on-site at -20 °C or in a dry shipper loaded with liquid nitrogen. Subsamples of the filtered fluids were reserved for major ion and trace element analysis in 50 mL acid washed falcon tubes. Samples for trace metal analysis were acidified with trace metal analysis grade nitric acid at 2% final concentration. Fluid aliquots were stored in 12 mL acid washed gas-tight vials for δ^13^C measurements of the dissolved inorganic carbon. Gas-phase and dissolved gas samples were collected in pre-evacuated 250 mL Giggenbach bottles containing 50 mL of 4 N NaOH to minimise atmospheric contamination^53^. In addition, gas-phase or dissolved noble gases were sampled at each site using copper tubes, as described previously^54^.

Sediment samples were collected at the fluid venting orifice using sterile, acid-washed PTFE spatulas and transferred into 50 mL Falcon tubes. Adjacent background soils were also sampled as controls (when present) from areas with no apparent geothermal influence, typically upsloping from the sample feature. These samples are used to track the surrounding soil microbial community contribution to the seeps fluid and sediments^18^. Background soil communities have no overlap with the community present in seeping fluids, and have minimum overlap with the sediment community. This, together with a suite of parameters previously described^18^ (and used in similar studies^9,8,5,11,55,56^), confirms the subsurface nature of the sampled fluids. All sediment samples were immediately frozen at -20 °C and kept at this temperature during transport to the laboratory, where they were stored until further analyses. Biofilm samples were collected using 10 mL sterile syringes or a spatula, and stored in individual 50 mL Falcon tubes, kept at -20 °C during transport to the laboratory until DNA extraction. The complete sampling approach^18^, Standard Operating Procedure^52^, and analytical methods^57^ have been described previously.

Alongside sample collection, physicochemical measurements of the fluids were performed *in situ*. Temperature was measured directly at the main vent outlet using a waterproof thermocouple. pH, oxidation-reduction potential, conductivity, and dissolved oxygen were measured with a multi-parameter probe (HANNA, HI98196) directly in the outlet or on fresh fluid samples kept in a headspace-less sealed container to allow for fluid cooling. pH, dissolved oxygen concentrations and saturation, and salinity were corrected for the difference in temperature between the *in situ* pool and the in-bottle quenched temperature (typically 50 °C) in case of seeps with temperatures above 55 °C. A fresh aliquot of filtered fluids was used in the field for the determination of redox or temperature sensitive parameters (see below).

### Geochemical measurements

In the field, total alkalinity, dissolved silica, and sulfide were analysed immediately after collection. Total alkalinity was determined by certified HCl titration (0.11%) using bromophenol blue as pH indicator to a terminal pH of 4.3 and calculated as mg of CaCO_3_ equivalent L^−1^ (HANNA, HI3811 Total Alkalinity Test Kit). Dissolved silica was determined using the HANNA HI97770C portable spectrophotometer and expressed as mg L^−1^ of SiO_2_. Total sulfide, expressed as mg L^−1^ of S^2-^ was determined using the HACH 1900 portable spectrophotometer and using the Cline assay.

Further geochemical measurements were performed on the samples in the laboratory. Ionic composition of fluids was determined by ion-exchange chromatography (IC) using a Metrohm ECO IC system, following the standardised procedures detailed previously^58^. To maintain conductivity below 600 μS/cm, which is critical for optimal chromatographic resolution, samples were diluted with 18.2 MΩ/cm type I water (typically 1:10). Cation and anion analyses were performed in separate runs. Anions were analysed using a Metrosep A Supp 5 column with a 3.2 mM Na_2_CO_3_ + 1 mM NaHCO_3_ mobile phase and a 0.15 M orthophosphoric acid suppressor. The anionic eluent was run at a flow rate of 0.7 mL min^−1^ for 30 min. Cations were analysed on a Metrosep C4 column using a 2.5 mM HNO_3_ plus 0.5 mM (COOH)_2_ × 2H_2_O mobile phase. The cationic eluent was run at 0.9 mL min^−1^ with a total separation time of 35 min. Data was acquired and analysed using MagIC Net 3.3 software. Calibration curves were generated using certified CPA Chem external standards for each anion and cation. Detection limits, defined at the 95% confidence level, were as follows: Mg^2+^ (0.08 mM), Na^+^ (0.3 mM), Ca^2+^ (0.3 mM), K^+^ (0.7 mM), SO_4_ ^2-^(0.03 mM), Br^−^ (0.02 mM), and Cl^−^ (0.1 mM).

Trace metal concentrations were measured by inductively coupled plasma mass spectrometry (ICP-MS) following the method described by ^59^. Briefly, fluid samples were acidified with HNO_3_ and diluted as a function of their salinity, while sediment samples were dried, homogenized, and digested using microwave-assisted acid digestion following the EPA3051A method. Analyses were performed on an Agilent 7900 ICP–MS. Calibration was carried out using multi-element standards spanning 0.01-100 µg L^−1^, with internal standards (e.g., Rh, In) added to correct for instrumental drift and matrix effects. Accuracy and precision were assessed using certified reference materials and quality-control standards, ensuring reliable quantification across a wide range of salinities.

Gas composition of gas-phase and dissolved gas samples, previously collected in Giggenbach bottles, was performed at the Observatorio Vulcanológico y Sismológico de Costa Rica (OVSICORI) at Universidad Nacional, Costa Rica, by gas chromatography (GC). Briefly, headspace gases (He, H_2_, O_2_, Ar, N_2_, and CH_4_) were quantified using an Agilent 7890a equipped with two HP-molesieve columns (Agilent 19095P-MSO) maintained at 30°C. Methane was analysed with a flame ionization detector while the other gases were analysed on thermal conductivity detectors. The NaOH solution from the Giggenbach bottles was then extracted and measured for CO_2_ content by titration with 0.1N HCl, allowing the calculation of the CO_2_/CH_4_ ratio based on the total number of moles of each gas. Carbon isotope compositions (δ^13^C) were determined in the NaOH solutions after acidification using a Picarro G2201-I analyser. The δ^13^C values, expressed in ‰ relative to the PDB standard, were calibrated on eight reference standards ranging from +2.42 ‰ to −37.21 ‰, including internationally recognised standards such as NBS19 and Carrara marble.

Noble gas analyses were conducted at the Barry Lab (WHOI) using a Nu Instruments Noblesse HR multi-collector mass spectrometer, capable of resolving all stable noble gas isotopes and separating key interferences (e.g., Ar- and CO_2_-based species) as previously described^54^. Copper tube samples (gas or water) were connected to a vacuum extraction line and expanded into a purification system. For water samples (∼13 mL), dissolved gases were first extracted under vacuum with magnetic stirring and quantitatively transferred using a capillary cryogenic method, whereby water vapour was trapped at −196 °C, entraining noble gases into a cold trap. Reactive gases were removed using a titanium getter at 650 °C, followed by hydrogen gettering during cooling and further purification through hot and cold SAES getters. Noble gases were then separated cryogenically: He and Ne were adsorbed on a charcoal trap (10–30 K), while heavier noble gases were retained on a stainless-steel trap. Helium and neon were sequentially released by stepwise heating and introduced into the mass spectrometer for isotopic analysis. Air standards were analysed routinely to monitor accuracy and instrument linearity, and procedural blanks were consistently <5% of the sample size. Interference corrections (e.g., ^40^Ar^++^, CO_2_ ^++^) were monitored but deemed negligible due to mass resolution and low background levels.

### DNA extraction and sequencing

DNA extraction from sediments and biofilms was performed either using the DNeasy PowerSoil Kit (QIAGEN) or a modified phenol-chloroform DNA extraction method adapted for low biomass samples^60^. Briefly, 0.8 g of sediment was suspended in 850 μL of extraction buffer (100 mM Na_2_HPO_4_ pH 8, 100 mM Tris–HCl pH 8, 100 mM EDTA pH 8, 1.5 M NaCl, 1% CTAB), supplemented with 100 μL lysozyme (100 mg/mL), and followed by two periods of incubation at 37 °C for 30 min, the second after the supplementation of 5 μL of Proteinase K (1 mg/mL). Then, 50 μL of 20% SDS was added and incubated at 65 °C for 1 h, with periodic mixing. After two steps of centrifugation performed at 14,000 g, the supernatant was collected and an equal volume of phenol:chloroform:isoamyl alcohol (25:24:1) was added DNA was supplemented with Na-acetate (0.1 vol) and isopropanol (100 %), incubated at room temperature for 12 h. After precipitation, DNA was washed with 70% ice cold ethanol, and finally resuspended in 50 μL Tris–HCl (50 mM). For selected low biomass samples, multiple extractions were combined. This phenol-chloroform extraction method was slightly modified to extract DNA from Sterivex filters 0.22 μm membrane, used to filter fluid samples^8^. Extracted DNA was subsequently sequenced by Novogene (UK) up to 100 million reads per sample using the NGS DNA Library Prep Set (Cat No.PT004) and Illumina platform NovaSeq X Plus. Before sequencing DNA samples were quantified using a Qubit (Invitrogen) dsDNA Broad Range or High Sensitivity assay and visually inspected on a 1% agarose gel coloured with EtBr.

### Public metagenomic datasets and associated environmental data

In addition to in-house produced metagenomes, we incorporated publicly available metagenomes from several sources. 110 publicly available subsurface metagenomes from public archives (NCBI SRA, DDBJ, ENA) were included. For surface comparison, 164 marine metagenomes were retrieved from the Tara Oceans^22,61^ (136 metagenomes, size fractions 0.22-1.6 and 0.22-3µm) and Malaspina^20^ (28 metagenomes, size fraction 0.2-0.8µm) expeditions. Soil and freshwater metagenomes were obtained from the National Ecological Observatory Network (NEON); soil metagenomes were collected between 2018 and 2021 (DP1.10107.001^62^), benthic freshwater metagenomes between 2018 and 2021 (DP1.20279.001^63^), and surface water metagenomes between 2014 and 2022 (DP1.20281.001^64^), yielding a total of 1,183 metagenomes before filtering. Associated environmental data were retrieved from the NEON soil physical and chemical properties product (DP1.10086.001^65^), the chemical properties of surface water product (DP1.20093.001^66^), and the periphyton, seston, and phytoplankton collection product (DP1.20166.001^67^). An additional 44 publicly available soil and freshwater metagenomes from individual studies were included. A list of all the metagenomes included in the analysis is available in Supplementary Data 1.

### Functional annotation and data filtering

In-house and collected published metagenomes were all analysed through the same pipeline. Raw reads were preprocessed using fastp^68^ set to default parameters allowing for trimming of low-quality bases, and functionally annotated using funprofiler (ksize = 11, scaled = 500, threshold_bp =500)^69^. Funprofiler is a pipeline leveraging FracMinHash sketching (implemented in Sourmash^70^) to functionally profile metagenomic samples against Kyoto Encyclopedia of Genes and Genomes (KEGG) database^71^. Intersect_bp, the estimated number of base pairs in common between the metagenome and the reference sketch of each KO, was taken as a metric of abundance of each KO within each metagenome. This metric was normalised by sample by the estimated abundance of the beta subunit of the RNA polymerase, chosen as a universal single-copy gene. *rpoB* is widely used as a single-copy, universal housekeeping marker across bacteria and archaea^72,73^, providing a proxy for genome equivalents and allowing comparison of functional gene abundance across samples. Practically, rpoB abundance was estimated by summing the normalised abundances of its different structural forms across Bacteria and Archaea (K03043 and K13797 in Bacteria; K13798 and the mean of K03044 and K03045 in Archaea), accounting for mutually exclusive gene configurations. This composite value was used as the normalisation factor for all KO abundances. For carbon fixation pathway abundance quantification, a set of markers was defined for each pathway (see Supplementary Note 1 and Supplementary Table 1), and the average abundance of this set was chosen as a proxy of pathway abundance in each metagenome.

To exclude low-quality samples, metagenomes from the CoEvolve, NEON, NCBI, and ENA datasets were filtered based on total rpoB abundance. Samples falling below the 5th percentile of the rpoB abundance distribution within each dataset were removed. Tara Oceans and Malaspina metagenomes were not subject to this filtering step as they had been previously quality-controlled. Additionally, samples in which fewer than 20% of KOs had non-zero abundances were removed from all datasets as low-annotation quality samples. Finally, only metagenomes with associated environmental data were retained (NEON soil samples being further filtered to those with the most complete metadata, proxied by the availability of δ^15^N measurements, to reduce their number in an unbiased way).

This resulted in a total 1,513 selected metagenomes including (i) 333 samples collected in the frame of the CoEvolve project (3 soil and 330 subsurface samples, of which 88 have been already published in previous work^5,74^), (ii) 82 subsurface and 41 surface metagenomes from public archives; (iii) 136 Tara Oceans and 28 Malaspina marine metagenomes, (iv) 376 and 517 soil and freshwater metagenomes from NEON. Altogether, these samples span a broad geographic and environmental space, and are compiled in Supplementary Data 1, with sample accession, project, and doi numbers indicated when relevant.

### Community composition and beta diversity

Beta diversity of rpoB-normalised KO abundance profiles was assessed using Bray-Curtis dissimilarity matrices computed with SciPy. Dimensionality reduction of the full dissimilarity matrix was performed using Uniform Manifold Approximation and Projection (UMAP). To test for significant differences in community composition among environment types (subsurface, marine water, freshwater, soil), PERMANOVA (permutational multivariate analysis of variance) was performed on the Bray-Curtis dissimilarity matrix with 999 permutations using scikit-bio. Homogeneity of multivariate dispersions among groups was assessed with PERMDISP (999 permutations). Principal Coordinates Analysis (PCoA) was applied to the Bray-Curtis dissimilarity matrix, and the distance of each sample to its environment group centroid was computed as the Euclidean distance from the sample’s PCoA coordinates (across all positive-eigenvalue axes) to the mean coordinates of its group, to visualize within-group compositional spread.

### Broad functional category analysis

For each metagenome, rpoB-normalised KO abundances were summed within each broad KEGG functional category and divided by the total summed category signal to get relative contributions of each functional category. Note that because some KOs are annotated to multiple functional categories, they were included in all relevant categories; therefore, category values are not mutually exclusive. KOs associated with carbon fixation pathways were reclassified into a dedicated “Carbon fixation” category to separate them from general metabolism for visualization purposes. The same approach was applied using the curated set of carbon fixation marker genes as categories to obtain a more targeted signal of carbon fixation pathway representation across samples. Stacked bar plots were generated to visualize the relative distributions of functional categories and carbon fixation marker genes across individual samples and environmental groups in two separate panels.

### Carbon fixation pathway analysis

Shannon diversity of carbon fixation pathway marker gene abundances was computed for each metagenome using scipy.stats.entropy. Differences in Shannon diversity between surface and subsurface environments were tested using the Mann-Whitney U test (scipy.stats.mannwhitneyu). For per-pathway comparisons of marker gene abundances between environment types, Mann-Whitney U tests were performed independently for each of the six carbon fixation pathways, with p-values corrected for multiple testing using the Holm-Bonferroni method (statsmodels). To visualize differential abundance of individual KOs between surface and subsurface environments, log_2_ fold-change was computed for each KO as log_2_(mean subsurface abundance / mean surface abundance) and plotted against mean log_10_ abundance across both environments (MA plot).

### Multivariate analysis of pathway-environment relationships

A principal component analysis of the carbon fixation pathway abundances was performed with scikit-learn. Environmental vector fitting was done using SciPy, with vector significance evaluated by permutation test (999 permutations) and FDR correction (Benjamini–Hochberg) with statsmodels. Spearman correlations between pathway abundances and environmental variables were calculated with SciPy and FDR-corrected with statsmodels.

Random forest models were trained independently for each carbon fixation pathway, using six environmental variables with high coverage across our subsurface samples dataset (Temperature, pH, DO, specific conductance, SO_4_^2-^, alkalinity; specific conductance, SO_4_^2-^, and alkalinity were log_10_-transformed prior to modelling) as predictors. 300 trees were built per model, with a square root feature sampling at each split, and a minimum leaf size of 5 (scikit-learn). The performance of each model was assessed by 5-fold cross-validated R^2^ and the statistical significance was evaluated using permutation testing (999 permutations; complete cases, n = 238). Variable importance was quantified as the mean decrease in hold-out R^2^ upon permutation of each variable (10 repeats per fold).

### Network analysis

Gene networks were built on carbon fixation genes only, using R, and the igraph package. These gene abundances underwent an ALR transformation (log-ratio against rpoB) to mitigate compositional effects, and genes with null-variance were removed. Spearman correlations were computed for each pair across all samples. Networks were built keeping only correlations above a threshold ρ, with edge weights equal to the correlation coefficient. Isolated nodes (degree = 0) were not kept.

In order to select an appropriate and unbiased ρ threshold, networks were built for ρ values from 0.30 to 0.80 (with 0.02 steps). Topological metrics were calculated at each step of this ρ-sweep. These included the node count, edge count, density, connected components, giant component fraction, mean degree, transitivity, and Louvain modularity. A ρ threshold of 0.45 was retained, corresponding to the value at which network remained connected, and modularity was close to its maximum. At this threshold, the community structure was identified using the Louvain algorithm. Modules containing fewer than three genes were removed. The stability of the different modules was tested by bootstrap resampling (200 iterations, 80% of samples each), recording gene co-assignment frequencies across iterations.

Two complementary metrics were additionally computed for each module, from rpoB-normalised abundances: (i) Latent module scores defined as the first principal component (PC1) of z-scored gene abundances within each module; (ii) Cumulated abundance of each module, defined as the sum of the rpoB-normalised abundances of all the genes of the module. The association of these two metrics with environmental variables was tested using Spearman’s correlation, restricted to variables with ≥ 20 observations, and with Benjamini–Hochberg FDR correction applied within each module. The same method was applied identically to both surface and subsurface datasets for consistency.

### Carbon Fixation Potential Index (CFPI)

For each carbon fixation pathway, pathway-level scores were calculated as the mean normalised abundance across their representative marker genes, normalised against the *rpoB* gene (see Functional annotation and data filtering). A Carbon Fixation Potential Index (CFPI) was defined as the sum of pathway-level scores and represents a relative proxy for autotrophic carbon fixation potential of each metagenome. Because *rpoB* approximates a single-copy reference and the occurrence of multiple carbon fixation pathways within a single genome is rare^2^, CFPI can be interpreted as an estimate of the fraction of the microbial community encoding carbon fixation potential. Using the mean abundance across pathway-specific marker genes minimises biases arising from differences in gene copy number and pathway architecture, ensuring comparability among pathways with different numbers of diagnostic genes.

Associations between CFPI and environmental variables were assessed using Spearman’s rank correlation with Benjamini–Hochberg correction (significance: p < 0.05, |ρ| ≥ 0.3). PCA was applied to standardised environmental variables to identify dominant geochemical gradients.

### Estimation of habitable crustal volume

Depth to the 135 °C isotherm (Z_135_) was calculated for each sample using a conductive geothermal gradient:

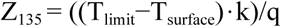

where T_limit_ = 135 °C, T_surface_ = 15 °C, k = 2.5 W m^−1^ K^−1^, and q is heat flow (W m^−2^). Heat flow values were obtained from a global gridded dataset^40^, using the predicted heat flow field to ensure continuous spatial coverage. Conductive thermal profiles in low heat-flow regions, Z_135_, were constrained by domain-specific maximum depths based on current knowledge of subsurface biosphere maximum depths. Oceanic settings were capped at 3 km, reflecting the upper permeable basaltic crust, whereas continental settings were capped at 10 km to approximate the depth limit of accessible, fracture-supported fluid circulation.

Pore-water volume was calculated for each grid cell as:

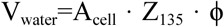

where A_cell_ is the grid cell area and ϕ is effective porosity. Porosity was assigned using literature-derived bulk estimates for the deep subsurface, with values of 5% for continental crust and 3% for oceanic crust, and sensitivity bounds of 3–8% and 1–5%, respectively, consistent with previous global estimates of subsurface pore space^41–43^. These values represent effective, connected porosity relevant to fluid circulation and microbial activity rather than total rock matrix porosity. Global habitable rock and pore-water volumes were obtained by summing across all grid cells and aggregating by continental and oceanic domains.

### Estimation of global carbon fixation flux

Since absolute rates cannot be inferred from gene abundances alone, CFPI was calibrated against published measurements from subsurface samples spanning different energy regimes and residence times. We used three published rate measurements as calibration anchors. The lower bound came from crystalline bedrock groundwater^45^, representing carbon fixation under strongly energy-limited conditions. For the intermediate anchor we used rates from carbonate aquifers^4^ which represent more active but still diffusion-limited subsurface fluids. The upper bound combined estimates from basalt-hosted systems^44^ and deep aquifers^46^, leaving out biofilm and near-surface measurements whose rates would not be representative of the subsurface. CFPI values coming from fluid-only samples were then mapped to this rate space (µg C L^−1^ d^−1^) through a piecewise log-linear interpolation that anchored the 5th, 50th, and 95th percentiles of the CFPI distribution to the low, intermediate, and high rate bounds respectively. Values outside this range were conservatively bounded. This approach effectively returns a metagenomic-informed statistical distribution of expected carbon fixation rates (r) across our global dataset that was used to determine the minimum, average and maximum carbon fixation values to use in the calculations. This calculation is an approximation based on the likelihood that the rate of carbon fixation is proportional to the relative abundance of chemolithoautotrophs in the microbial community. Total rate was estimated by the formula:

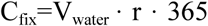

and expressed in Pg C yr^−1^. Uncertainty ranges were propagated from both CFPI variability and porosity bounds.

## Data availability

Raw sequences are available through the European Nucleotide Archive (ENA) under the Umbrella Project CoEvolve PRJEB55081. All samples used in this study, together with their associated geochemical and environmental data, are compiled in Supplementary Data 1. For publicly available metagenomes, accession numbers, project identifiers, and DOIs are indicated when relevant.

## Code availability

Code and metadata containing all the steps to reproduce our analysis and figures are available at https://github.com/giovannellilab/Subcarb-Global_organization_of_carbon_fixation released in Zenodo with DOI 10.5281/zenodo.20053398.

## Acknowledgments

BdP was supported by the Marie Skłodowska-Curie Actions Postdoctoral Fellowship (Grant Agreement No. 101154017; HORIZON-MSCA-2023-PF-01 project SUBCARB) under the European Union’s Horizon Europe programme. This work was supported by funding from the European Research Council (ERC) under the European Union’s Horizon 2020 research and innovation program Grant Agreement No. 948972—COEVOLVE—ERC-2020-STG to DG. Funding was also provided by the European Union’s Horizon 2020 research and innovation programme (Grant Agreement No. 948972, COEVOLVE, ERC-2020-STG) and by the European Union’s Clean Hydrogen Joint Undertaking (Grant Agreement No. 101192337, HYDRA). PHB, KGL, and MdM acknowledge NSF award 2121637, and PHB and KGL acknowledge NSF 2152551, which partially supported this work. Additional support came from The National Fund for Scientific and Technological Development of Chile (FONDECYT) Grant 1251543 (The National Research and Development Agency of Chile, ANID Chile), COPAS COASTAL ANID FB210021 to GLJ. DG and AVB were supported by the Italy MUR PRIN2022 grant (Grant No. 20224YR3AZ; HYDECARB). DG and JB were also supported by the European Union’s Horizon 2020 research and innovation programme (Grant Agreement No. 871120; International Network for Terrestrial Research and Monitoring in the Arctic, INTERACT; Transnational Access call, GHOST project). JAB was also supported by the Agence Nationale de la Recherche (ANR23-CPJ1-0172-01) and the European Research Council (ERC) under the European Union’s Horizon Europe Research and Innovation programme (Grant agreement No. 101115755, acronym SIESTA). GCG was supported by the European Union’s Horizon Europe Research and Innovation programme under the Marie Skłodowska-Curie Grant Agreement No. 101073271, project SHINE. We thank Secretaría de Medio Ambiente de la Provincia de Catamarca for their administrative support that allowed our sampling (Expte.EX-2022-02222431-CAT-DPB#SEAS, Resolución S.E.A., D.S. N°: 053/2017), the Secretaría de Ambiente y Desarrollo Sustentable – Ministerio de Producción y Desarrollo Sustentable de la provincia de Salta (IF-2025-62727469-APN-SSAM#JGM, Resolución N°315/25), the Secretaría de Biodiversidad y Desarrollo Sustentable – Ministerio de Ambiente y Cambio Climático de la provincia de Jujuy (IF-2025-50987272-APN-SSAM#JGM, Resolución SBDS N°061/25), the Ministry of Business, trade, Mineral Resources, Justice and Gender Equality of Greenland (license no. G24-026), the Environment and Energy Agency of Iceland (license n. UST202305-096/D.Ö.H. and OS2023010131/50.4.3), the Department of Environment and Tourism of Uvurkhangai aimag of Mongolia (permit n. 144 of 2023.07.26). Additionally, we acknowledge Secretaría de Política Ambiental en Recursos Naturales, Ministerio de Ambiente y Desarrollo Sostenible de la Nación Argentina for providing the certificate of compliance (Title: IF-2023-65966642-APN-SPARN#MAD, UId: ABSCH-IRCC-AR-264943-1) and the export certificate for genetic resources (CE-2023-69835755-APN-SPARN#MAD). We thank Lucas Paoli and Elad Noor for their constructive feedback on this work. We are also grateful to the Arctic Station in Qeqertarsuaq for logistical support in the field.

## Competing interests

The authors declare no competing interests.

## CoEvolve Project Consortia

Donato Giovannelli, CoEvolve consortium PI, Department of Biology, University of Naples “Federico II”, Naples, Italy

Angelina Cordone, Department of Biology, University of Naples “Federico II”, Naples, Italy

Agostina Chiodi, CONICET Instituto de Bio y Geociencias del NOA (IBIGEO), Rosario de Lerma, Argentina

Alberto Vitale Brovarone, Dipartimento di Scienze Biologiche, Geologiche e Ambientali -BiGeA, Università degli Studi di Bologna Alma Mater, Bologna, Italy

Alessia Bastianoni, Department of Biology, University of Naples “Federico II”, Naples, Italy

Annarita Ricciardelli, Department of Biology, University of Naples “Federico II”, Naples, Italy

Bernardo Barosa, Department of Biology, University of Naples “Federico II”, Naples, Italy

Carlos Ramirez Umana, Servicio Geológico Ambiental (SeGeoAm), Heredia, Costa Rica

Davide Corso, Department of Biology, University of Naples “Federico II”, Naples, Italy

Deborah Bastoni, Department of Biology, University of Naples “Federico II”, Naples, Italy

Edoardo Taccaliti, Department of Biology, University of Naples “Federico II”, Naples, Italy

Federico A. Vignale, European Molecular Biology Laboratory -Hamburg Unit, Hamburg, Germany

Feliciana Oliva, Department of Biology, University of Naples “Federico II”, Naples, Italy

Flavia Migliaccio, Department of Biology, University of Naples “Federico II”, Naples, Italy

Francesco Montemagno, Department of Biology, University of Naples “Federico II”, Naples, Italy

Gabriella Gallo, Department of Biology, University of Naples “Federico II”, Naples, Italy

Gerdhard L. Jessen, Instituto de Ciencias Marinas y Limnológicas, Universidad Austral de Chile, Valdivia, Chile

J. Maarten de Moor, Observatorio Vulcanológico y Sismológico de Costa Rica (OVSICORI), Universidad Nacional, Heredia, Costa Rica

Jacopo Brusca, Department of Biology, University of Naples “Federico II”, Naples, Italy

James A. Bradley, Aix Marseille Univ, Université de Toulon, CNRS, IRD, MIO, Marseille, France

Karen G. Lloyd, Earth Science Department, University of Southern California, Los Angeles, CA, USA

Luca Tonietti, Department of Science and Technology, Parthenope University of Naples, Naples, Italy

Luciano di Iorio, Department of Biology, University of Naples “Federico II”, Naples, Italy

Maria García Alai, European Molecular Biology Laboratory -Hamburg Unit, Hamburg, Germany

Martina Cascone, Department of Biology, University of Naples “Federico II”, Naples, Italy

Matteo Selci, Department of Biology, University of Naples “Federico II”, Naples, Italy

Mattia Esposito, Department of Biology, University of Naples “Federico II”, Naples, Italy

Monica Correggia, Department of Biology, University of Naples “Federico II”, Naples, Italy

Mustafa Yucel, Institute of Marine Sciences, Middle East Technical University, Erdemli, Turkey

Nunzia Nappi, Department of Biology, University of Naples “Federico II”, Naples, Italy

Peter H. Barry, Marine Chemistry & Geochemistry Department, Woods Hole Oceanographic Institution, MA, USA

Sara Claudia Diana, Department of Biology, University of Naples “Federico II”, Naples, Italy

Stefano Caliro, Osservatorio Vesuviano, Istituto Nazionale di Geofisica e Vulcanologia (INGV), Napoli, Italy

## Notes

https://github.com/giovannellilab/Subcarb-Global_organization_of_carbon_fixation

### Competing Interest Statement

The authors have declared no competing interest.

