## Supplementary Information for "Global carbon fixation in Earth’s subsurface biosphere"

#### Contents

**Supplementary Notes** **2**
Supplementary Note 1. 2
Supplementary Note 2. 2

**Supplementary Tables** **4**
Supplementary Table 1. 4

**Supplementary Figures** **6**
Supplementary Fig. 1. 6
Supplementary Fig. 2. 6
Supplementary Fig. 3. 7
Supplementary Fig. 4. 7
Supplementary Fig. 5. 8
Supplementary Fig. 6. 9
Supplementary Fig. 7. 10
Supplementary Fig. 8. 11
Supplementary Fig. 9. 12
Supplementary Fig. 10. 13
Supplementary Fig. 11. 14

### Supplementary Notes

#### Supplementary Note 1. Detecting carbon fixation enrichment requires pathway-specific marker genes

The detection of carbon fixation pathways enrichment in the subsurface requires the establishment of a list of marker genes for each pathway. Indeed, considering the entire set of genes involved in each pathway dilutes the signal, since most of them are shared with other pathways, often participating in central metabolism. To illustrate this effect, we progressively examined the functional annotation of metagenomes at increasing levels of specificity, from broad functional categories to individual carbon fixation pathways, and finally to marker genes specific to these pathways (Supplementary Fig. 4 and Supplementary Table 1 for the list of selected marker genes). At the level of global functions, metabolism, or energy metabolism (including, specifically, carbon fixation pathways), we do not observe any significant enrichment of subsurface gene abundance, compared to surface (soil, fresh and marine waters). However, when limiting the analysis to marker genes specifically associated to carbon fixation pathways only, a clear enrichment is observed for subsurface samples (Supplementary Fig. 4, lower part outlined with a dashed line). This approach isolates genes that are exclusive to these pathways, and therefore give a more precise representation of the carbon fixation potential of each metagenome. More generally, these results show the importance of defining sets of marker genes that are specific and exclusive to metabolic pathways for functional analysis. By focusing on them rather than on the full set of pathway-associated genes, such approaches reduce dilution of the signal, and therefore allow a more precise detection of ecological and metabolic profiles across different environments.

#### Supplementary Note 2. Controlling the effect of sequencing-depth on the observed signal

Since the current study uses datasets of different sources, and since the studied KOs can be relatively rare in the metagenomes, we sought to assess whether the results discussed in this paper could result from a sequencing bias rather than a biological reality. Indeed, even if KO abundances are divided by abundance of the beta subunit of the RNA polymerase, chosen as a universal single-copy gene, that would technically normalise by cell count, one can not exclude the possibility that a signal can emerge solely from whether detection thresholds are crossed at certain sequencing depths. Especially, since most subsurface metagenomes studied here originate from our lab and were sequenced to high depth (up to 100M reads per sample using Illumina NextSeq technology), we can indeed observe significantly higher sequencing depths (median read count one order of magnitude higher) in subsurface versus surface metagenomes (Supplementary Fig. 6a). Since the observed higher abundance of carbon fixation pathways in the subsurface (Fig. 2 and Supplementary Fig. 5) might result from such a technical bias, we therefore re-performed the analysis with subgroups of surface and subsurface metagenomes sharing the same median read count. Briefly, for each subsurface metagenome, we matched the nearest neighbour based on the read count, compared on a logarithmic scale with a tolerance of 0.2 (meaning a factor of ×1.6 between the read counts within paired subsurface and surface metagenomes). These two matched sets of 345 subsurface and surface metagenomes each exhibit a similar median read count as shown in Supplementary Fig. 6b. The higher abundance of carbon fixation pathways observed in Fig. 2 are maintained in these subgroups (Supplementary Fig. 6c), confirming that the effect is not a sequencing-depth threshold artifact but rather describes a biological reality.

### Supplementary Tables

**Supplementary Table 1.** KEGG Orthology (KO) marker genes diagnostic of the six carbon fixation pathways investigated in this study.

| Pathway | KO | Gene name | Gene description | EC number |
|---|---|---|---|---|
| **Calvin-Benson-Bassham (CBB)** | K00855 | PRK/prkB | phosphoribulokinase | 2.7.1.19 |
| | K01601 | rbcL/cbbL | ribulose-bisphosphate carboxylase large chain | 4.1.1.39 |
| | K01602 | rbcS/cbbS | ribulose-bisphosphate carboxylase small chain | 4.1.1.39 |
| **3-Hydroxypropionate bicycle (3HP)** | K09709 | meh | 3-methylfumaryl-CoA hydratase | 4.2.1.153 |
| | K14468 | mcr | malonyl-CoA reductase / 3-hydroxypropionate dehydrogenase (NADP+) | 1.2.1.75 |
| | K14469 | K14469 | acrylyl-CoA reductase (NADPH) / 3-hydroxypropionyl-CoA dehydratase / 3-hydroxypropionyl-CoA synthetase | 1.3.1.84 |
| | K14470 | mct | 2-methylfumaryl-CoA isomerase | 5.4.1.3 |
| | K14471 | smtA1 | succinyl-CoA:(S)-malate CoA-transferase subunit A | 2.8.3.22 |
| | K14472 | smtB | succinyl-CoA:(S)-malate CoA-transferase subunit B | 2.8.3.22 |
| **3-Hydroxypropionate/ 4-Hydroxybutyrate (HPHB)** | K14466 | K14466 | 4-hydroxybutyrate---CoA ligase (AMP-forming) | 6.2.1.40 |
| | K15018 | K15018 | 3-hydroxypropionyl-coenzyme A synthetase | 6.2.1.36 |
| | K15019 | K15019 | 3-hydroxypropionyl-coenzyme A dehydratase | 4.2.1.116 |
| | K15020 | K15020 | acryloyl-coenzyme A reductase | 1.3.1.84 |
| | K15039 | K15039 | 3-hydroxypropionate dehydrogenase (NADP+) | 1.1.1.298 |
| **Dicarboxylate/ 4-Hydroxybutyrate (DCHB)** | K14467 | 4hbl | 4-hydroxybutyrate---CoA ligase (AMP-forming) | 6.2.1.40 |
| **Wood-Ljungdahl (WL)** | K00192 | cdhA | anaerobic carbon-monoxide dehydrogenase/CODH/ACS complex subunit alpha | 1.2.7.4 |
| | K00193 | cdhC | acetyl-CoA decarbonylase/synthase/CODH/ACS complex subunit beta | 2.3.1.169 |
| | K00194 | cdhD/acsD | acetyl-CoA decarbonylase/synthase/CODH/ACS complex subunit delta | 2.1.1.245 |
| | K00195 | cdhB | anaerobic carbon-monoxide dehydrogenase/CODH/ACS complex subunit epsilon | |
| | K00197 | cdhE/acsC | acetyl-CoA decarbonylase/synthase/CODH/ACS complex subunit gamma | 2.1.1.245 |
| | K00198 | cooS/acsA | anaerobic carbon-monoxide dehydrogenase catalytic subunit | 1.2.7.4 |
| | K14138 | acsB | acetyl-CoA synthase | 2.3.1.169 |

| | | | | |
|---|---|---|---|---|
| **Reductive TCA (rTCA)** | K15230 | aclA | ATP-citrate lyase alpha-subunit | 2.3.3.8 |
| | K15231 | aclB | ATP-citrate lyase beta-subunit | 2.3.3.8 |
| | K15232 | ccsA | citryl-CoA synthetase large subunit | 6.2.1.18 |
| | K15233 | ccsB | citryl-CoA synthetase small subunit | |
| | K15234 | ccl | citryl-CoA lyase | 4.1.3.34 |
| | K00176 | korD/oorD | 2-oxoglutarate ferredoxin oxidoreductase subunit delta | 1.2.7.3 |
| | K00177 | korC/oorC | 2-oxoglutarate ferredoxin oxidoreductase subunit gamma | 1.2.7.3 |

1

### Supplementary Figures

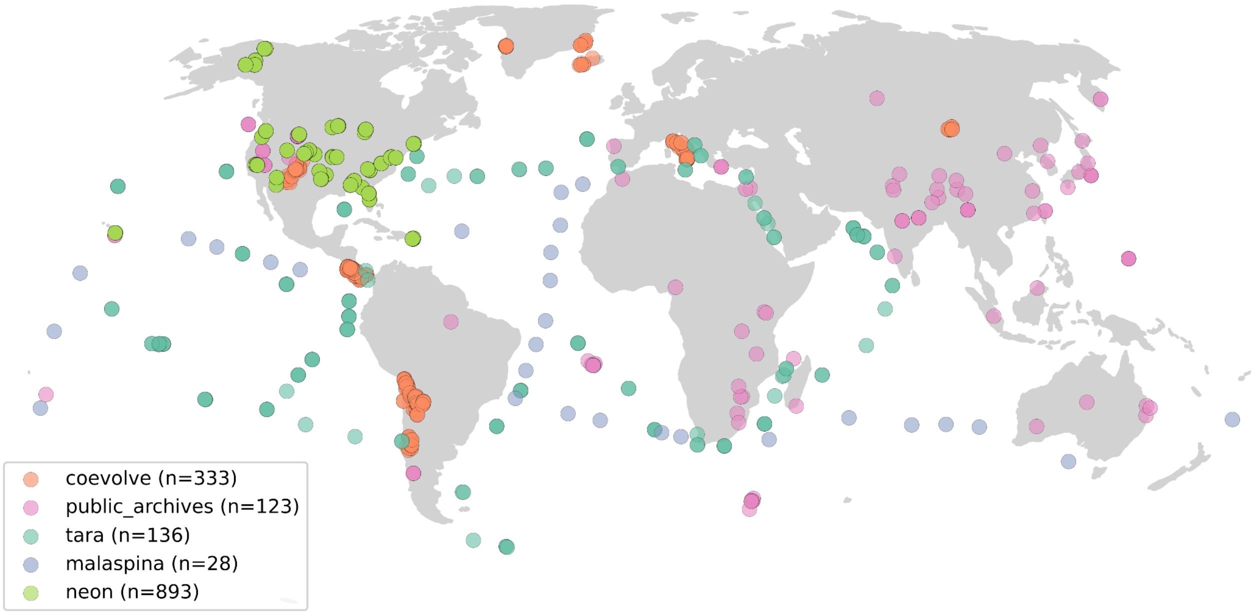


**Supplementary Fig. 1: Global distribution of subsurface and surface metagenomes shown by data source.**
Samples included in this study comprise metagenomes from the CoEvolve dataset, together with surface marine (Tara Oceans, Malaspina), freshwater and soil (Neon), and additional subsurface and surface metagenomes from public archives (NCBI SRA, DDBJ, ENA).

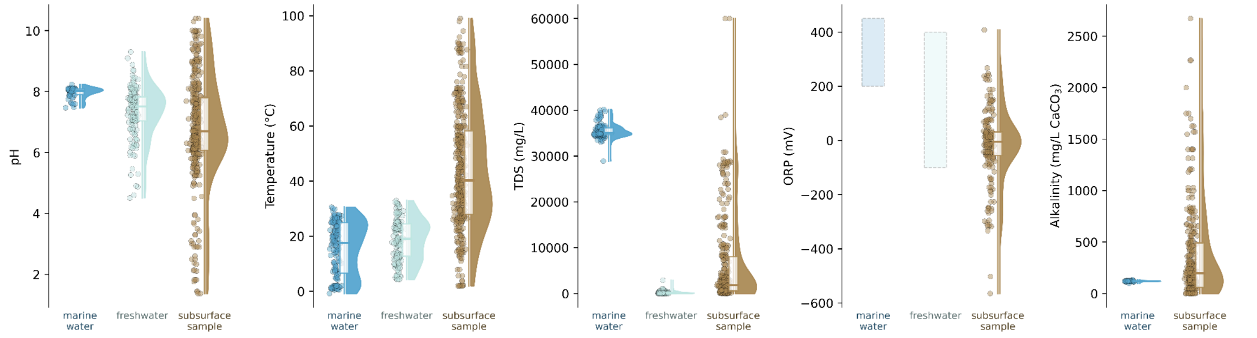


**Supplementary Fig. 2: Physicochemical distributions of subsurface fluids, freshwater, and marine samples across environments.**
Subsurface fluids harbour higher ranges of pH, temperature, total dissolved solids (TDS), oxidation–reduction potential (ORP), and alkalinity. Points represent individual samples; half-violins show kernel density distributions; boxes indicate interquartile ranges with median lines. Dashed rectangles in the ORP panel indicate literature ranges for freshwater and marine waters (Stumm & Morgan 1996; Millero 2013; Appelo & Postma 2005).

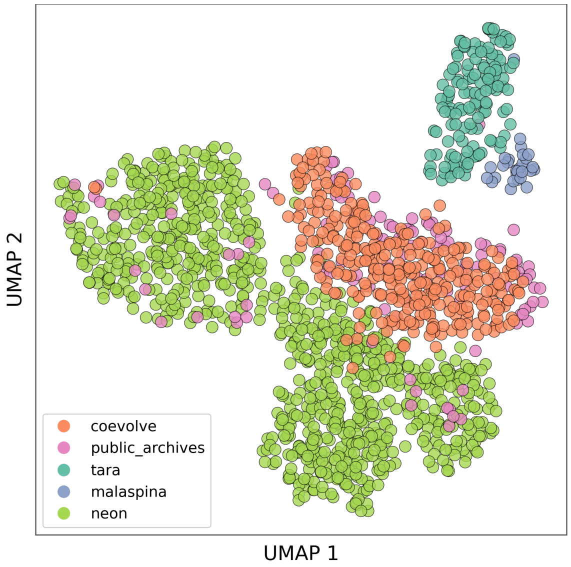


**Supplementary Fig. 3: UMAP of all metagenomic samples, based on functional profiles, coloured by sample source.**

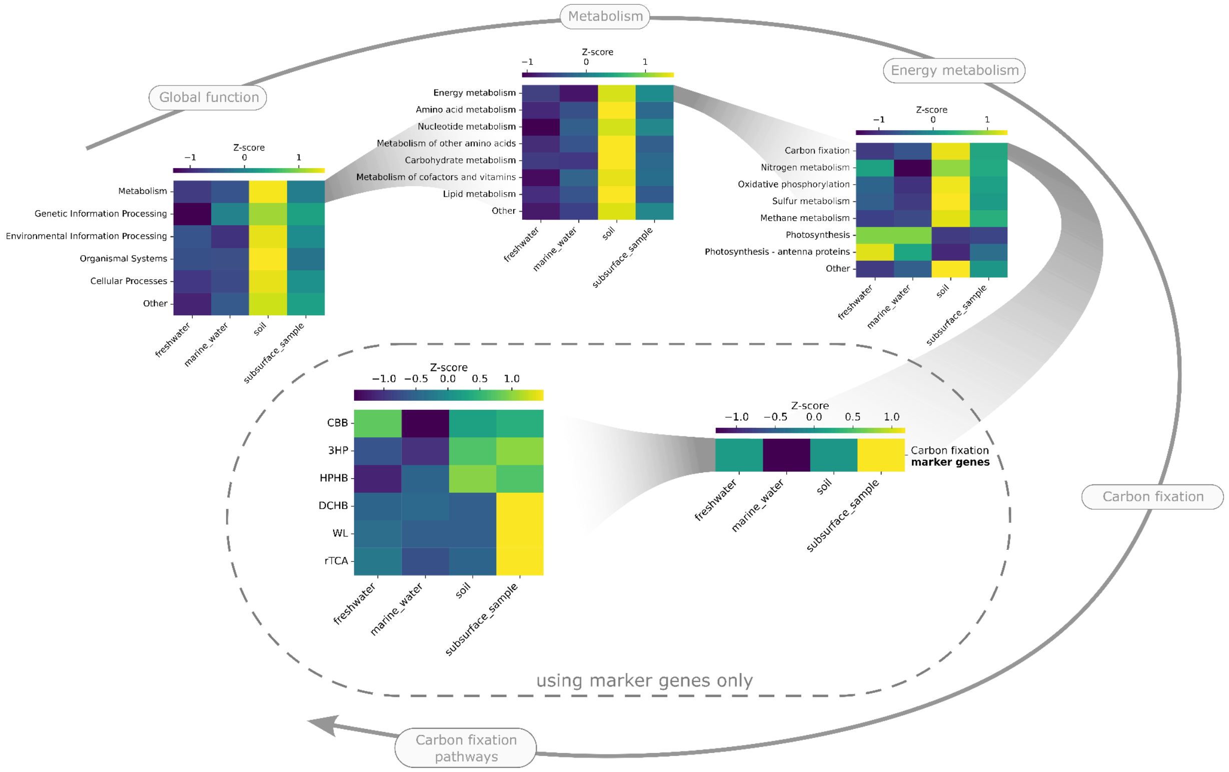


**Supplementary Fig. 4. Carbon fixation enrichment in subsurface metagenomes is only detectable using pathway-specific marker genes.**

KO abundances were averaged by function across sample types at different KEGG pathway hierarchies. Within each functional category, values were normalised using row-wise z-scores to show relative enrichment patterns.

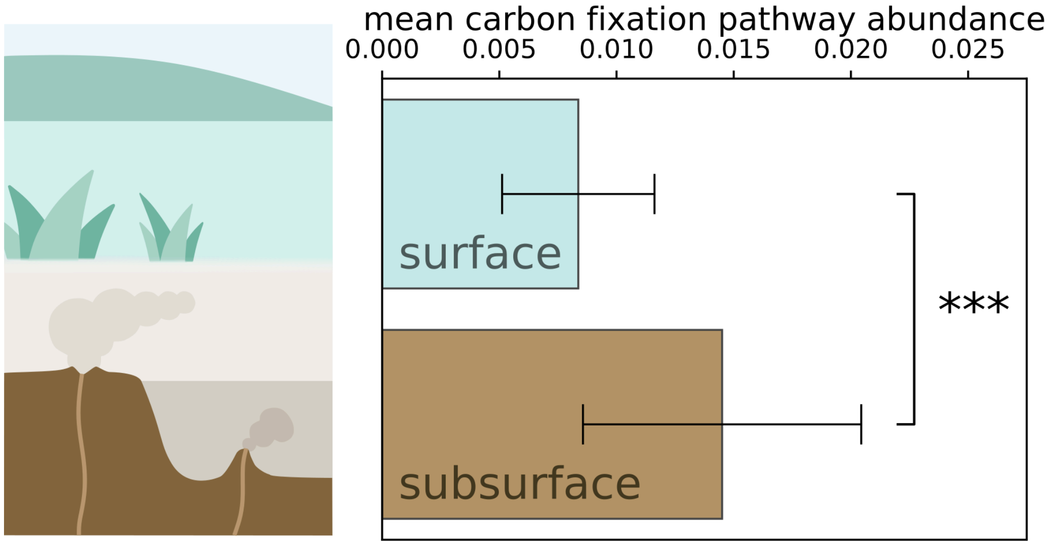


**Supplementary Fig. 5. Subsurface enrichment of carbon fixation genes compared to the surface.** Average abundance of carbon fixation marker genes in the surface and subsurface metagenomes of this study (Mann-Whitney U test, *** $p < 0.001$).

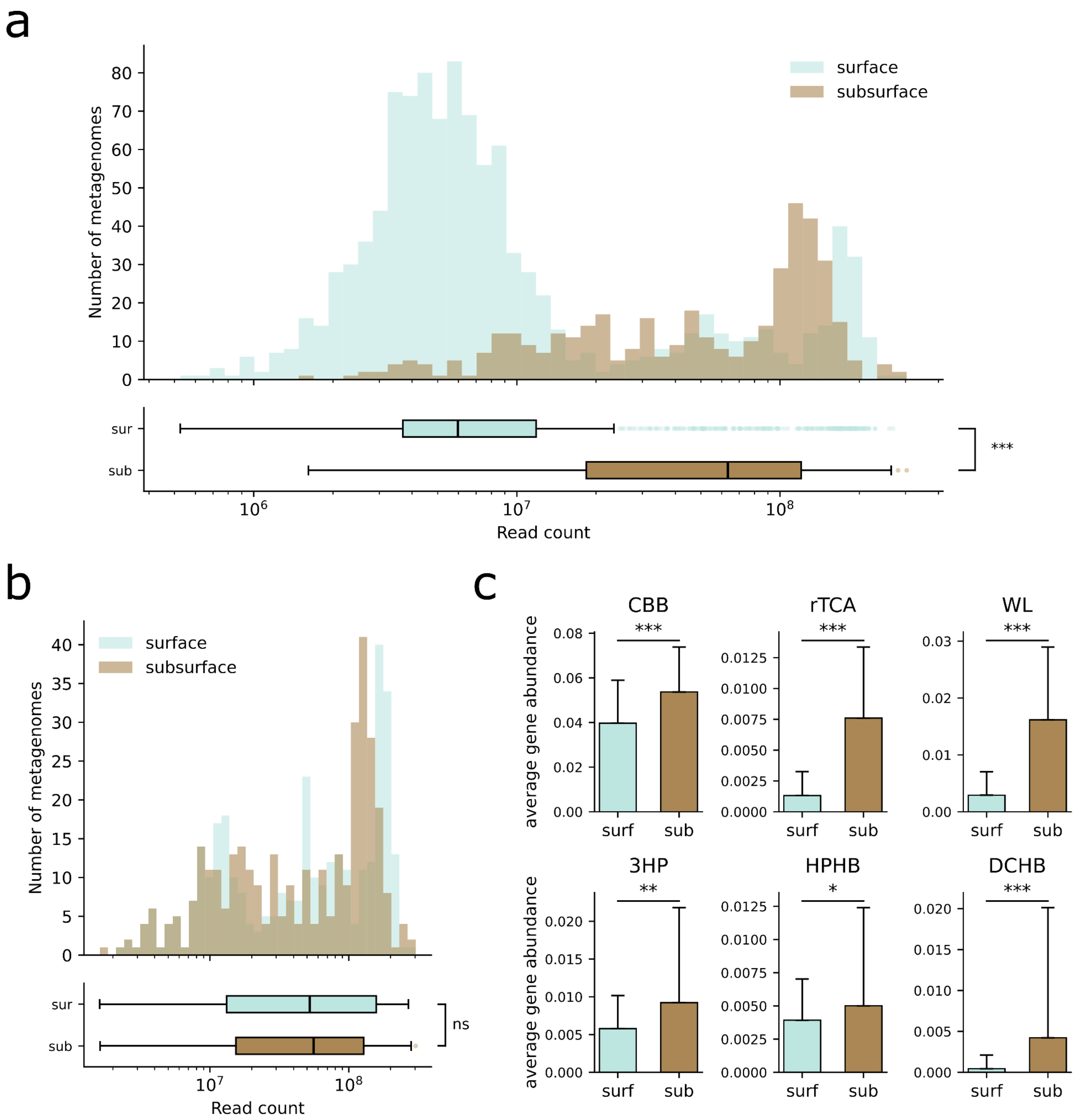


**Supplementary Fig. 6. Carbon fixation genes abundance enrichment in subsurface metagenomes does not result from a sequencing depth artifact.**

**a,** Distribution of total read counts in surface (light blue) and subsurface (brown) metagenomes across the full dataset. Subsurface metagenomes show significantly higher sequencing depth than surface metagenomes (Mann-Whitney U test, $p < 0.001$). **b,** Same as a, but restricted to same-depth subsets built by nearest-neighbour matching on read counts (Mann-Whitney U test, ns: not significant). **c,** Mean gene abundance of carbon fixation pathway markers in the surface and subsurface subsets from b. Differences assessed by Mann-Whitney U test (* $p < 0.05$, *** $p < 0.001$, ns: not significant).

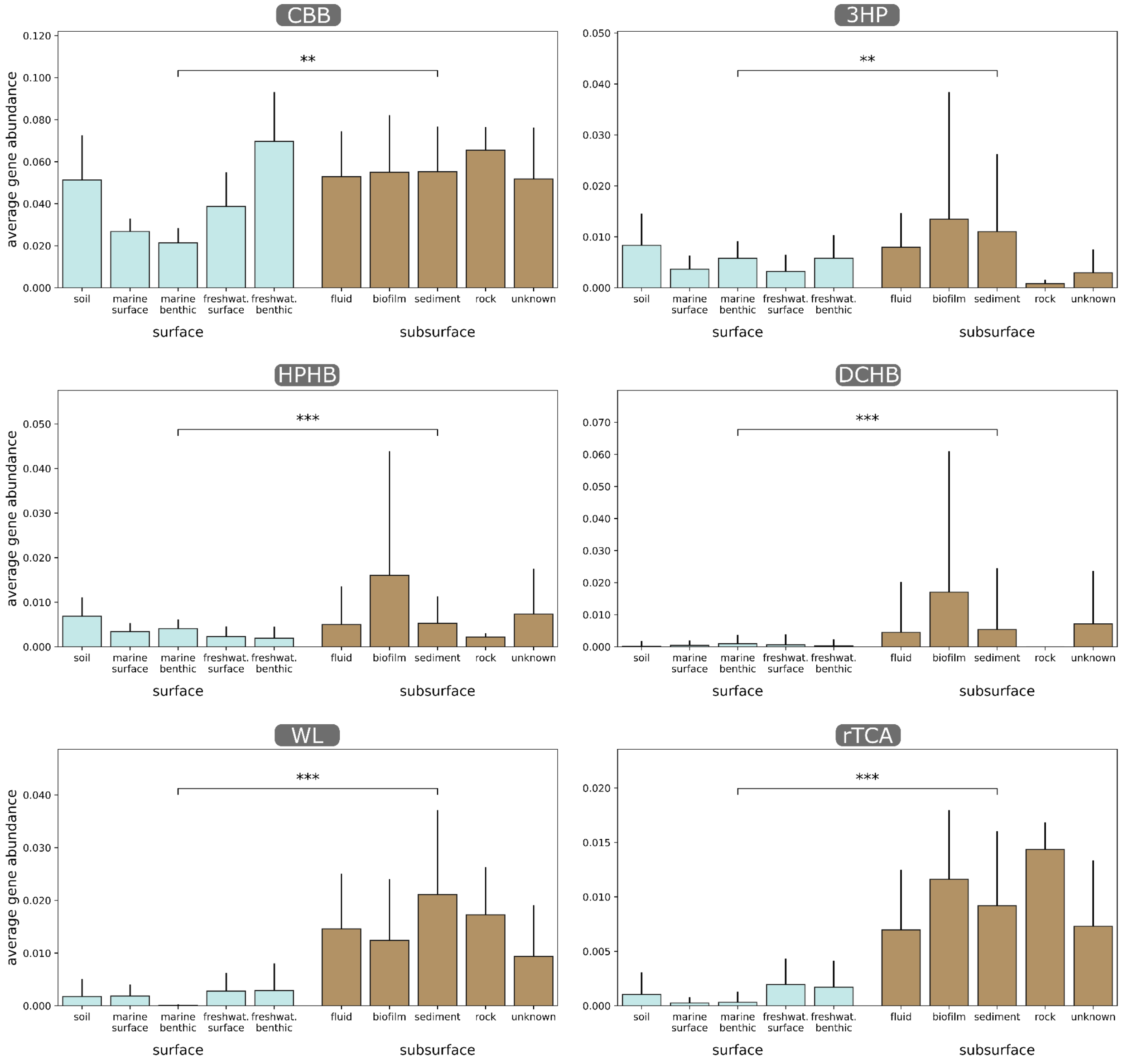


**Supplementary Fig. 7. Pathway-level comparison of carbon fixation strategies across all sample types.**

“Unknown” refers to subsurface samples retrieved from public repositories for which detailed sample type metadata was not available. Differences assessed by Mann-Whitney U test (** $p < 0.01$, *** $p < 0.001$).

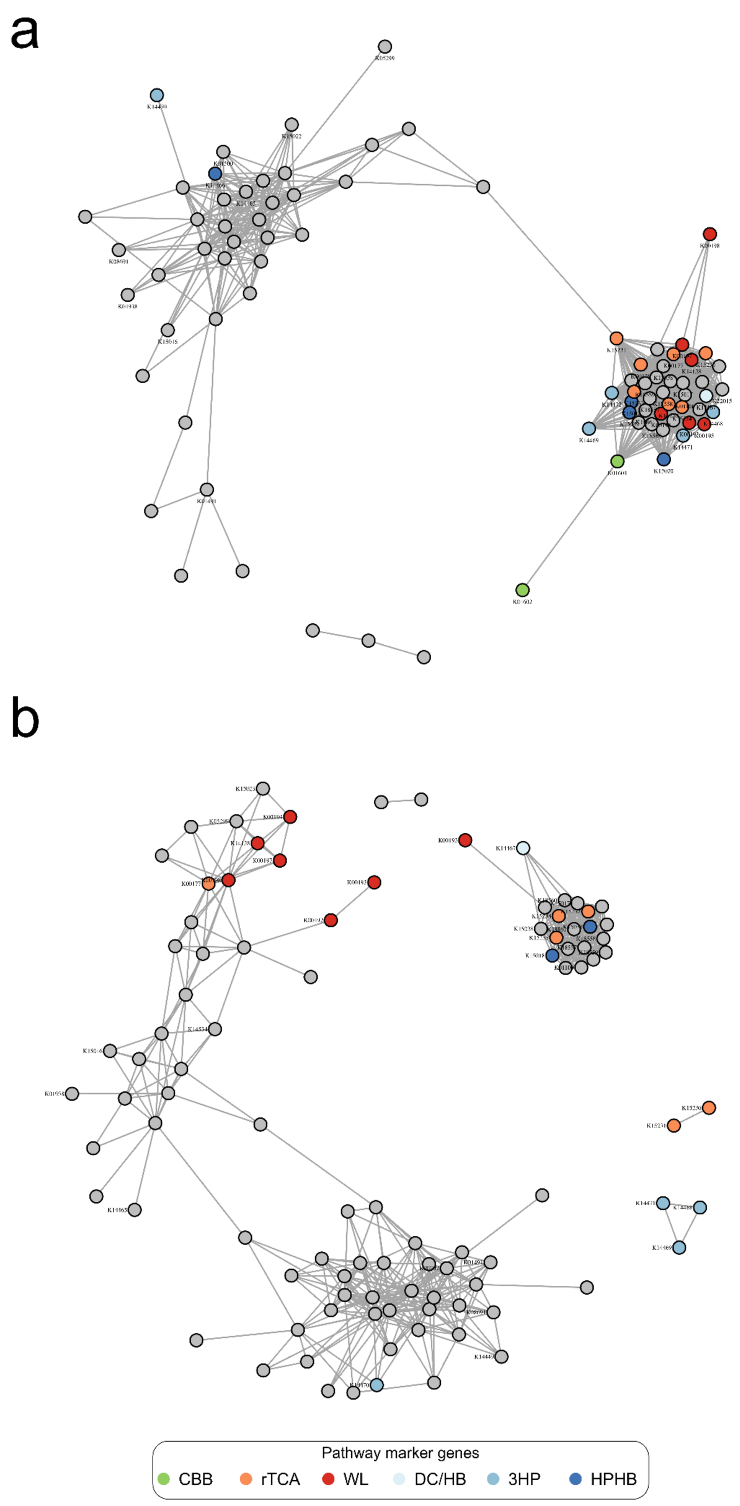


**Supplementary Fig. 8. Subsurface carbon fixation gene network exhibits higher modularity than the surface network.**

Surface (**a**) and subsurface (**b**) carbon fixation gene network topology at ρ=0.45. The surface network is composed of 83 nodes and 940 edges, with a modularity of 0.307. The subsurface network is composed of 96 nodes and 471 edges, with a modularity of 0.614.

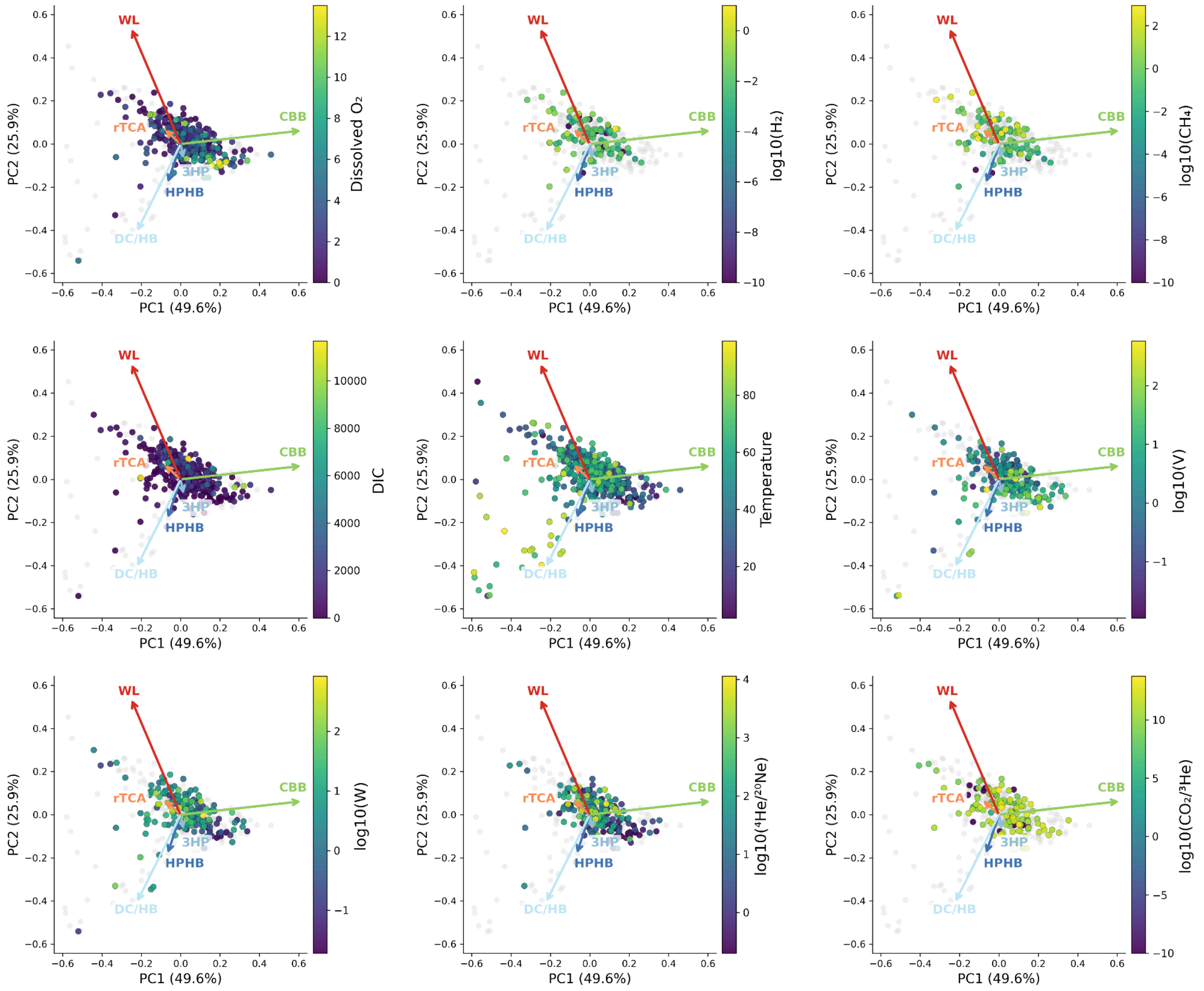


**Supplementary Fig. 9. Environmental gradients projected onto the carbon fixation pathway ordination in subsurface samples.**

Samples are shown in the PCA space of the six carbon fixation pathways (same ordination as Fig. 4c) and coloured according to selected environmental variables, including temperature, dissolved $O_2$, dissolved inorganic carbon (DIC), gas concentrations ($H_2$, $CH_4$), deep fluid tracers ($^4He/^{20}Ne$, $CO_2/^3He$), tungsten and vanadium.

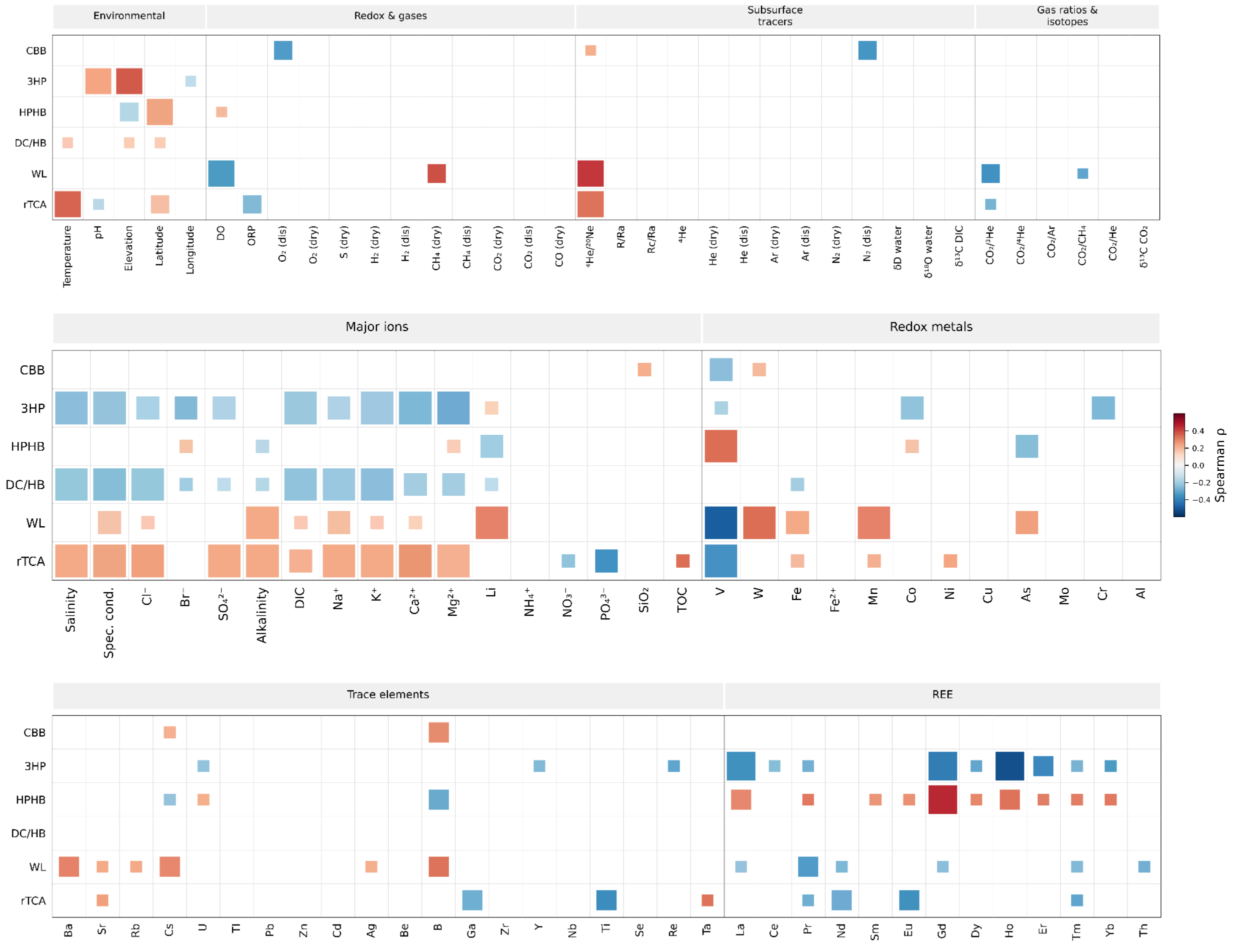


**Supplementary Fig. 10. Spearman correlations between carbon fixation pathways and environmental parameters measured in this study.**

Square size represents statistical significance (FDR-BH q-value) and colour indicates the correlation coefficient (ρ).

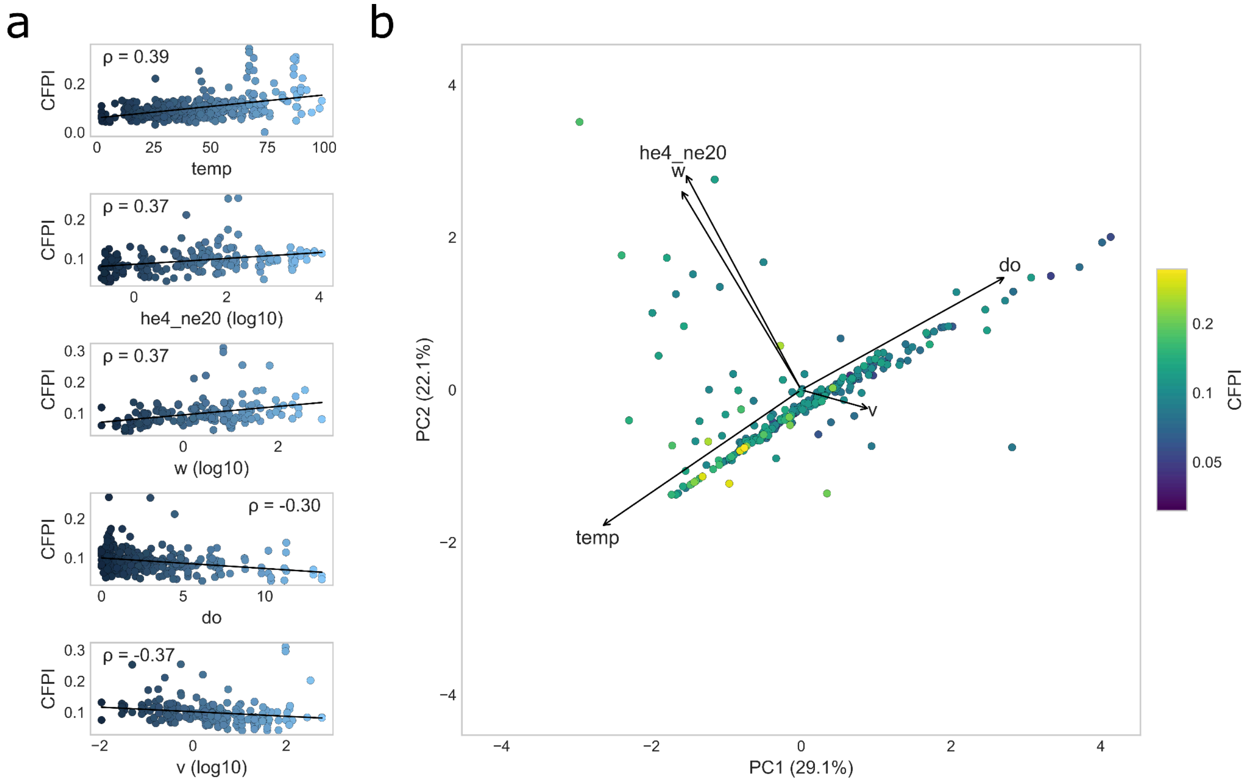


**Supplementary Fig. 11. Carbon Fixation Potential Index (CFPI) globally correlates along an axis linking redox state, temperature, and fluid residence time.**

**a,** CFPI plotted against temperature, $^{4}He/^{20}Ne$, tungsten, dissolved oxygen (do), and vanadium concentration. Spearman coefficients are indicated for each relationship. **b,** PCA of subsurface samples based on these five environmental parameters. Dots are coloured according to their CFPI. Environmental variables are represented as loading vectors.
